# Tradeoffs in bacterial physiology determine the efficiency of antibiotic killing

**DOI:** 10.1101/2022.03.09.483592

**Authors:** Anat Bren, David S. Glass, Yael Korem Kohanim, Avi Mayo, Uri Alon

## Abstract

Antibiotics can kill or stop the growth of bacteria, and their effectiveness depends on many factors. It is important to understand the relation between bacterial physiology, the environment and antibiotic action. While many of the mechanistic details of antibiotic action are known, the connection between death rate and bacterial physiology is poorly understood. Death rate in antibiotics has often been shown to rise linearly with growth rate; however, it remains unclear how environmental factors, in concert with whole-cell physiological properties, affect bactericidal activity. To address this, we developed a high-throughput assay to precisely measure antibiotic-mediated bacterial death. We found that death rate is linear in growth rate, but the slope depends on environmental conditions. Specifically, stressors lower the death rate compared to a non-stressed environment with the same growth rate. To understand the role of stress, we developed a mathematical model of bacterial death based on resource allocation that takes into account a newly defined stress-response sector; we identify this sector using RNA-seq. Our model accurately predicts the death rate and minimal inhibitory concentration of antibiotics across a wide range of conditions, including a previously unknown increase in the stress response and protection from death at very low levels of cAMP. The present death-growth model suggests conditions that may improve antibiotic efficacy.

## Introduction

The first antibiotic was discovered over 100 years ago ^1^. Since then, many antibiotics that either kill the bacteria (bactericidal) or primarily inhibit their growth (bacteriostatic) have been discovered ^2^. The direct interactions and proximal mechanisms of action have been elucidated for many antibiotics. However, the connection between the molecular mechanism of action and the physiological state of the bacterium (e.g., growth rate, proteomic profile) that ultimately leads to death remains poorly understood ^2–4^. Understanding how bacteria deal with antibiotics is particularly relevant due to increasing issues of resistance mutations^5–7^.

Another concern in antibiotic treatment is tolerance, a natural ability to survive prolonged treatment ^8^. Tolerance is not accompanied by a change in the minimal inhibitory concentration (MIC) ^8^ and is known to depend on the bacterial growth environment ^8–10^. Because of the clinical importance, many studies have approached antibiotic efficacy from the perspective of outcome (i.e., bacterial death) rather than the physiological state of the bacterium (see ^2^ for a recent review). In this paper, we focus on the connection between death rate, MIC and the physiological state of the bacteria.

Previous studies found linear relationships between growth rate or metabolic state and death rate due either to bactericidal antibiotic ^11–17^ or to starvation ^18^. In contrast to these simple linear relations, combinatorial treatments show greater complexity (see ^19^ for a recent review). For instance, bacteriostatic antibiotics protect against death due to bactericidal antibiotics ^20^ and anti-ribosomal antibiotics protect against anti-DNA antibiotic ^21^. In addition, starvation and other stressful conditions were found to protect bacteria from antibiotics ^9,10,22–25^. It is thus reasonable to expect that the connection between growth rate and death rate is more complex than a simple linear function of growth rate and may be dependent on the environment and the physiological state of the cell.

Unlike the death rate, the growth rate has been extensively studied in terms of physiology, revealing simple growth laws ^26–30^. An early example of a growth law is the linear increase of growth rate with ribosomal content ^27^ (the basis of an “R sector” in later terminology ^30^). This simple growth law has proven impressively capable of predicting bacterial growth rates across a wide range of environmental conditions, despite the thousands of underlying molecular reactions ^26,27,30^. More detailed yet still coarse-grained models extended the growth laws to predict growth rate as a function of multiple internal proteomic sectors, each representing large groups of genes with similar behavior under corresponding resource limitations ^31–34^. For example, the “C sector” represents genes which are upregulated under carbon limitation ^31,35,36^.

One attempt to relate resource allocation to death proposed that investment in maintenance prolongs survival during starvation ^18^. We hypothesized that a generalized resource-allocation model, which takes into account tradeoffs between sectors due to limited cell resources, could connect environmental conditions, internal bacterial physiology and antibiotic killing rates. To build and test such a model requires accurate measurement of death rates in many conditions.

In this paper, we developed a high-throughput method to measure bactericidal death rates in a variety of conditions. We found that the death rate does not depend on growth rate alone, but also on the details of the environment. Stressful environments protect against bactericidal killing relative to non-stressed environments with the same growth rate. We hypothesized that stressful environments activate a cellular physiological program that helps bacteria to deal with damage imposed by the antibiotic treatment ^37^. To test this, we developed a mathematical model that can quantitatively recapitulate death rates from given environmental conditions, based on tradeoffs in the allocation of resources to growth-related and stress-related proteomic sectors. Moreover, the model could accurately predict MIC, which we measured in an independent manner and which rose with decreased death rate only under stressful conditions – an effect we term *hardiness*. We confirmed the existence of such a stress sector using RNA sequencing and quantitatively validated the model predictions of C (carbon), S (stress), and R (ribosomal) sector sizes in various conditions. By directly manipulating the sector sizes via cAMP, we found a surprising decrease in death as well as an increase in MIC at low cAMP, which is quantitatively predicted by the model. Finally, we use our results to discuss the clinical relevance and suggest treatment conditions that may improve antibiotic killing of bacteria.

## Results

### High throughput assay of bacterial death rates

Bacterial death rates are classically measured via the colony-forming unit (CFU) assay. This assay estimates the number of viable bacteria remaining after various times in a damaging treatment by counting colonies that grow after plating on permissive agar media. This method is labor-intensive and limited in throughput. High-throughput measurements of decreasing optical density (OD) ^12^ or of minimal duration of killing (MDK) ^38^ are either limited to specific antibiotics that disrupt cell integrity (e.g., ampicillin) or yield limited time-course information. Single-cell tracking of death via microfluidics ^39^ is not easily scaled to measurement in many conditions. Overall, there is a lack of robust, high-throughput methods to measure death curves.

Here, we developed such an automated, high-throughput assay to measure death rates in 96-well plates on a robotic system (details in Fig. 1 and Methods). In short, we measured the surviving fraction of cells as a function of antibiotic challenge duration. The robotic system enabled us to run a reverse time course with antibiotic introduced into exponentially growing cultures at a consistent OD (Figs. 1A, S1). Following the start-growth-time method of Hazan, *et al*., we estimated surviving cell numbers by measuring the time τ for a treated culture to reach a certain OD threshold in permissive (minimal glucose) media (Fig. 1B)^40^. Fewer live cells growing exponentially will take longer to reach the threshold and thus represent lower percent survival, which we quantified by comparison to delays τ in untreated, diluted cultures (Fig. 1C). From percent survival in a range of antibiotic treatment durations, we obtained a survival curve and fit it to a Weibull survival function coupled to exponential growth (see Methods). Note that in some conditions we observed an initial increase in the number of viable bacteria, reflecting that bacteria at first grew faster than they died (Figs. S2,S3), as has been observed previously ^12^. We defined death rate as 1/t_90_, the inverse of time to reach 10% survival of the initial population (Figs. 1D, S3, Methods).

**Fig 1.**
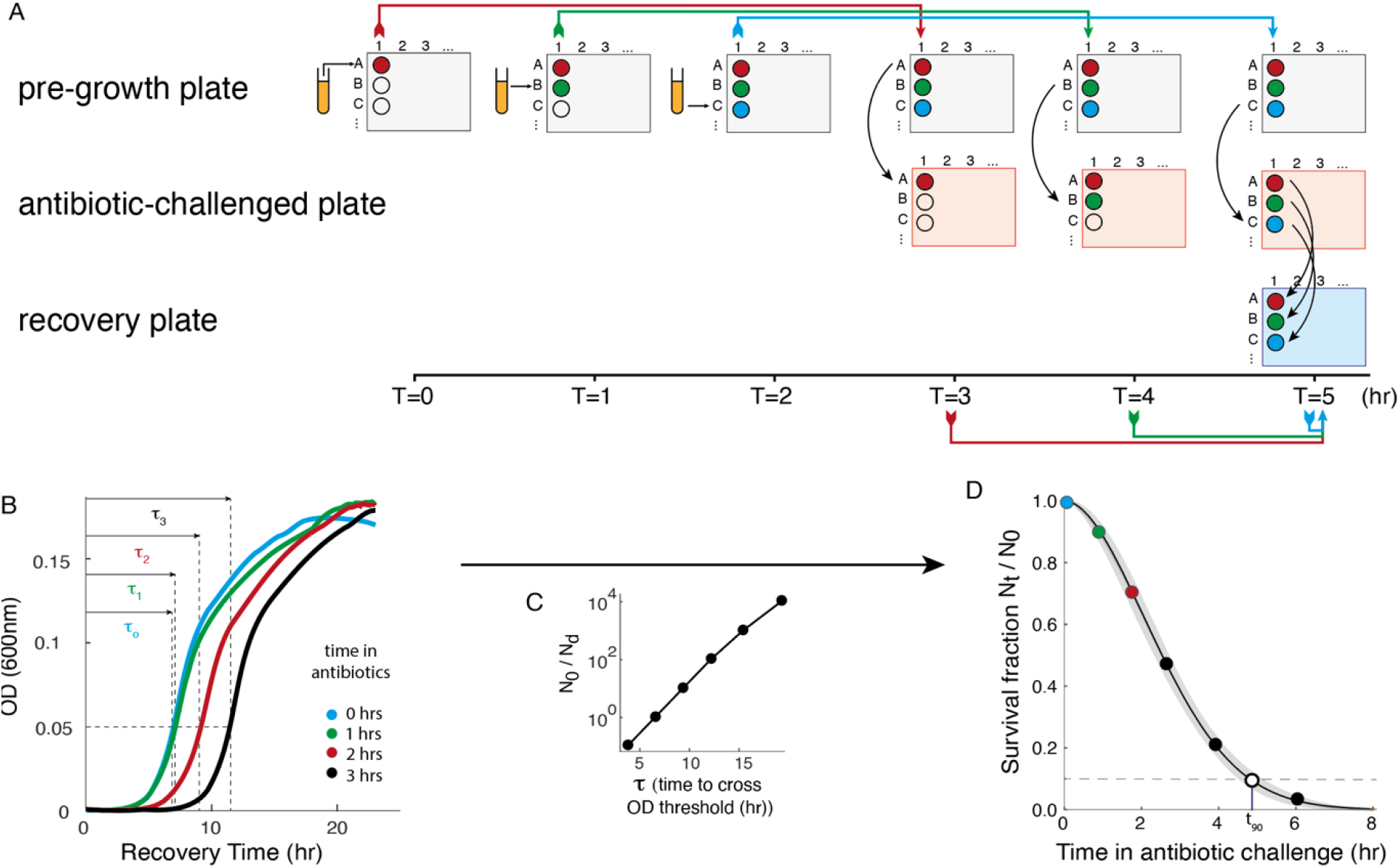
Death rate measurement protocol. For simplicity, we illustrate here the process for 3 time points (additional time points indicated in black) for one replicate of a single growth condition, with 1-hr resolution. **A**. Scheme of the robotic process. (i) *Pre-growth*: in this stage bacteria are inoculated each hour in successive wells. (ii) *Antibiotic challenge*: after 3 hours of pre-growth, each exponentially growing culture is transferred to antibiotics. (iii) *Recovery*: all cultures are moved to antibiotic-free recovery medium at the same time, resulting in cultures that have been treated with antibiotics for various time durations. (i.e., wells in different colors have the same pre-growth conditions but spend different times in antibiotic) **B**. OD curves of cultures recovering from the antibiotic treatment. For each curve, we define the delay time τ to cross the OD=0.05 threshold indicated by the horizontal dashed line. Cultures that spent more time in antibiotics have larger τ. **C**. A standard curve obtained from the τ values of non-treated cultures with a range of dilutions (*N*_0_ = initial concentration, *N*_*d*_ = diluted concentration) gives the relation between delay and relative number of bacteria. **D**. Surviving fraction as a function of time in the antibiotic challenge is calculated based on the measured τ and the standard curve (*N*_0_ = concentration of live cells at time 0, *N*_*t*_ = concentration of live cells at time t). We fit this data with a Weibull survival function (plus growth for initially growing cultured, Fig. S2-S3, main text) and defined death rate as 1/t_90_, where t_90_ is the time for the function to drop to 10%.

We validated this approach by comparison to the CFU method in various conditions and found very good agreement (Fig. S4, Methods). We also tested the sensitivity of the method to treatment duration and found that the calibration between recovery time τ and survival does not depend on time in antibiotic (Fig. S5). We conclude that the high throughput assay provides an accurate measure of killing in the present conditions.

Overall, the protocol provided throughput of 4-8 survival curves in a two-day experiment (see Figs. S2 and S15 for all death curves obtained in this study).

### Death rate depends on both growth rate and physiological stress

Using this assay, we explored the relation between growth and death rates of *E. coli* NCM3722 under various physiological conditions (Figs. S2, S3). As a challenge, we used 10μg/ml of the bactericidal antibiotic nalidixic acid, which interferes with DNA gyrase ^41^. We used multiple growth conditions and evaluated the growth rate in each in the absence of the antibiotic challenge. First, we studied the effect of various carbon sources in a minimal growth medium. We found, in agreement with previous studies, that the lower the growth rate, the lower the death rate. Death rate in glucose, which supported the fastest growth, was the highest; death rate was lowest in the poorest carbon sources, galactose and mannose (blue dots, Fig. 2)

**Fig 2.**
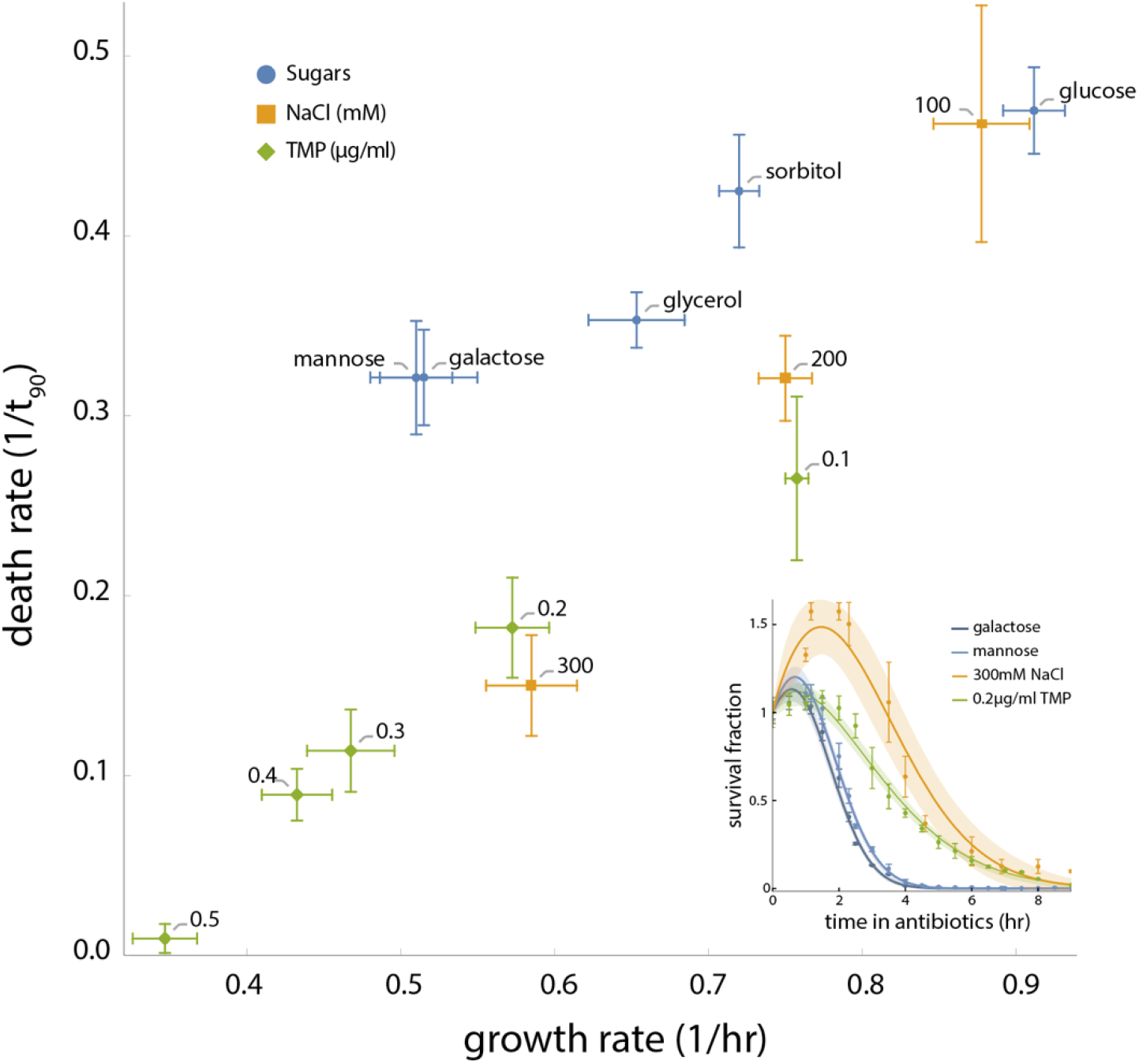
Antibiotic-mediated death depends on both growth rate and growth condition. The dependency of death rate on growth rate for NCM3722 strain (measured separately without nalidixic acid) under stress is steeper than under non-stressful conditions. Death rate upon treatment with 10 μg/ml nalidixic acid as a function of growth rate is shown for 13 different conditions (M9 + glucose as the reference point, M9 + 4 additional carbon sources and M9 + glucose + varying concentrations of NaCl or TMP). Each rate is determined based on at least 3 biological repeats. Error bars are standard error. **Inset**. At similar growth rates (0.52-0.59 hr^-1^), lower death rates in stress conditions (NaCl and TMP) derive from wider survival curves than present in non-stressed conditions (mannose and galactose). Shown are averaged survival curves (see also Fig. S3), with shaded areas the 95% confidence interval of the fit (Methods).

We next used glucose as a carbon source and reduced the growth rate by applying stress conditions, namely conditions that limit growth not by nutrients but through other environmental parameters^34^. Specifically, we used NaCl at high osmolality or the DNA-damaging antibiotic trimethoprim (TMP), which is bacteriostatic in minimal media^42,43^.

We found that the growth-death relation was steeper in stressors than in the carbon sources. In other words, we death rate depends on the environment, with stressful conditions providing dose-dependent protection (orange and green dots, Fig. 2). Further increase in stress levels (400mM and 500mM NaCl) resulted in cells growing faster than they died, which we quantified as a negative death rate (Fig S6, Methods).

Thus, conditions with a similar growth rate may result in significantly different death rates. For example, the death rate on mannose or galactose was ∼2-fold higher than the death rate reached by 300mM NaCl or 0.2μg/ml TMP, with much steeper survival curves (Fig. 2 inset). Protection from death was also found when using ethanol and tetracycline as stressors (Fig. S7). This finding was not exclusive to Nalidixic acid. We measured death rate by phosphomycin, an antibiotic from a different class (a membrane synthesis inhibitor^44^) and similarly found that death rate on mannose was higher than on glucose plus 300mM NaCl or 0.2μg/ml TMP (Fig S8). We also tested thye aminoglycoside antibiotic streptomycin^45^ and found protection from death by 300mM NaCl, but not by 0.2μg/ml TMP (Fig. S8).

We conclude that death rate is not solely a function of growth rate. Antibiotic causes a higher death rate when applied to bacteria growing on a poor carbon source than when it is applied to bacteria with the same growth rate growing on a rich carbon source supplemented with stressors.

### A resource allocation model can explain the observed growth-death data

We hypothesized that a resource-allocation model including a stress-response sector can explain the observed growth and death rates. We begin with the dependence of death rate on growth rate in different carbon sources, which we call the ‘sugar line’ (Fig. 3A, blue line). On the sugar line, the well-established resource allocation theory predicts that as more resources are dedicated to carbon catabolism (C sector), fewer resources are dedicated to building ribosomes (R sector), resulting in a correspondingly lower growth rate^31,35,36^. One may assume that under these conditions the death rate increases in direct correlation to growth rate due to increased production of damage in line with previous descriptions^11,15^ (e.g., nalidixic acid affects DNA gyrase, which introduces DNA breaks during replication^46^).

**Fig. 3.**
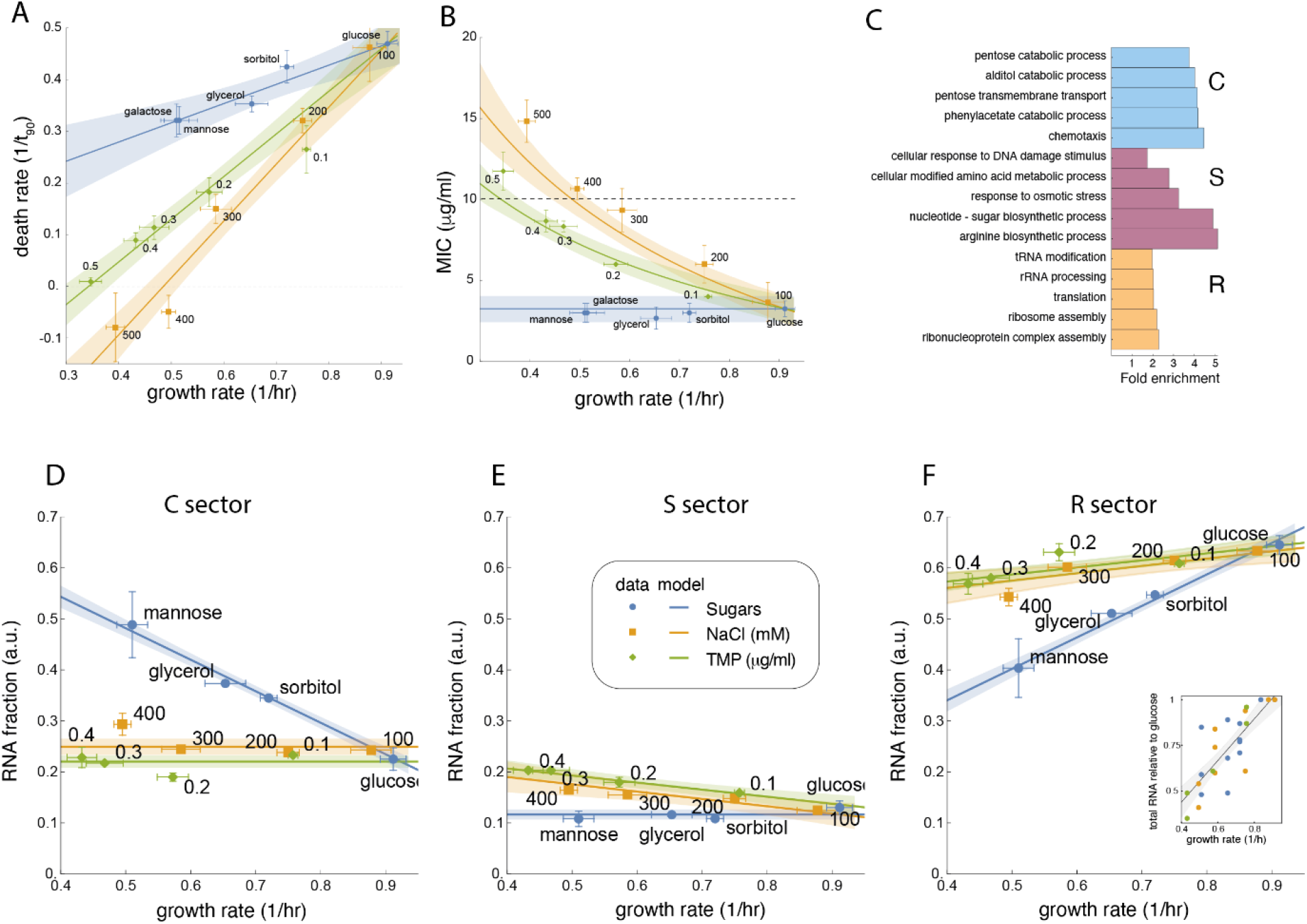
A resource-allocation model quantitatively matches growth, death, MIC, and sector-size measurements. **A**. Fitting Eqs 1-3 (Methods) to growth and death measurements from Fig. 2 produces a tight fit, with S constant for sugar data and C constant for stressors. **B**. The model predicts MIC data with no additional free parameters if the direct effect of growth on death is assumed to increase linearly with nalidixic acid concentration. **C**. Gene ontology (GO) terms for the determined C, S, and R sectors match expected behaviors of catabolism, stress, and growth, respectively. The stress sector contains anabolic processes, presumably due to indirect stress-response needs. Displayed terms are 5 terms with lowest false discovery rate that are not child-terms of other significant terms. **D-F**. Total mRNA fraction of the measured sectors show that C is relatively constant with increasing stress while S is relatively constant regardless of sugar quality. R increases with growth rate, but at different slopes for sugars and stressors. **F inset**. Total RNA increases with growth rate at the same slope for both sugars and stressors. Shaded regions in all panels other than **B** are 90% confidence intervals on the fitted parameters (Methods). For **B** the error bands are propagated from the fits in **A** (Methods). Legend for **A, B, E-F** as in **E**.

In contrast, when growth rate is varied via stressor concentration using glucose as a carbon source, we expect the C sector to remain constant. We base this expectation on the fact that the carbon source remains unchanged and the stresses imposed are unrelated to metabolic constraints. We predict that with stressors present, resources are redirected to a newly defined stress-response sector (S sector) at the expense of the R sector. This increase in stress-related genes can provide protection against antibiotic damage while decreasing growth rate, resulting in a ‘stress line’ (Fig. 3A, green and orange lines). We assume that under the conditions we studied, changing resources are divided strictly into the R, C and S sectors, while other sectors remain unchanged. This hypothesis predicts that conditions in which both carbon source and stress level change will yield growth and death rates that reside between the sugar and stress lines. Indeed, we find that combinations of glycerol+TMP and glycerol+NaCl lie between these lines, as does acetate, which is a poor carbon source known to induce a stress response ^47,48^ (Fig S9).

Based on the above hypotheses, we developed a mathematical resource-allocation model, summarized by the following equations. We use growth on glucose as an anchor point and define the change of the sectors in a given condition as ΔC, ΔS and ΔR. Since the sum of all sectors is constant (C+S+R=1), their total change must equal zero ^30^:

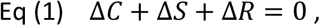

The growth rate *µ* depends linearly on the R sector as described by the well-established growth law ^27,30^ *µ* = *aR* − *b*, and thus the change of growth rate relative to glucose obeys:

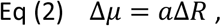

where *a* is the ribosomal growth efficiency.

The new aspect of the model is an equation for the death rate. Death rate ρ increases with damage, which is proportional to growth rate and reduced by the S sector, leading to the proposed death law:

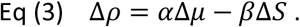

Here Δ*ρ* is death rate minus death rate on glucose, *α* is the decrease in death rate per decrease in growth rate and *β* is the death protection efficiency per unit increase in the S sector. We assume that on the sugar line, which lacks stressors, S remains constant so that Δ*S* = 0, while on the stress line C remains constant so that Δ*C* = 0 and thus 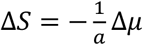. Fitting Equations 1-3 to the growth-death measurements (Fig. 3A, Table S1, Methods) provides an excellent fit (adj. R^2^= 0.986) with two non-dimensionalized free parameters, 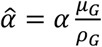 and 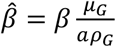 (Methods). The sugar line slope Provides 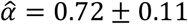. The stress line shows that protection by NaCl 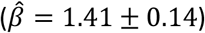 is greater than for TMP 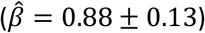.

### The resource allocation model accurately predicts MIC as a function of growth rate

We next considered the three major concepts of bacterial survival in antibiotics: resistance, tolerance and persistence, as recently defined by Brauner *et al* ^8^. Two are not relevant to this study – persistence relates to a very small subpopulation that is not killed, whereas our experiments focus on the whole-population level. Resistance is due to genetic changes, which do not occur in our short-term experiments. The remaining concept, tolerance, is defined as the ability of microorganisms to survive antibiotic treatment for a longer time (reduced death rate) without a change in MIC ^8^. We therefore set out to measure the MIC in order to test whether the reduced killing in our conditions is due to tolerance.

We quantified MIC by measuring growth curves in a range of nalidixic acid concentrations, identifying the MIC as the lowest antibiotic concentration that prevents growth (Methods, Fig. S10). We found that MIC does not vary with different sugars (Fig. 3B, blue points), corresponding to tolerance. However, MIC increased with stressors in a dose-dependent manner (Fig. 3B, green and orange points) with negative correlation to death rate (Fig. S11). This requires a new concept to describe reduced killing accompanied by increased MIC, which we term *hardiness*.

Indeed, both hardiness and tolerance are predicted quantitatively by our death model. Mathematically, we characterize MIC by *ρ* = 0, corresponding to a flat survival curve. Thus, we expect that conditions lying on the horizontal axis in Fig. 3A to have a MIC of 10μg/ml nalidixic acid (the concentration used). Assuming that the growth-dependent damage *α* and death rate on glucose *ρ*_*G*_ increase linearly with antibiotic concentration, we derive a relation between MIC and growth rate for all growth conditions (Methods). Specifically, MIC remains constant for the sugar line but increases according to a Michalis-Menten-like function of growth rate. Thus, the model predicts tolerance for sugars and hardiness for stressors. Without any additional free parameters, the model prediction provides an excellent fit to the MIC data (adj. R^2^=0.982, Fig. 3B).

### Gene-expression measurements support the prediction of a sizable stress sector

A basic assumption of the proposed model is a sizable S sector, whose fraction of cellular resources at high stress is significant enough to lead to a decrease in the R sector and a corresponding decrease in growth rate.

To examine the size and composition of the S sector, we performed RNA-Seq analysis on *E. coli* NCM3722 cultures grown in various carbon sources or in glucose with increasing concentrations of NaCl or TMP (Methods). We grouped genes into clusters using a Gaussian mixture model and then grouped the clusters by increasing, decreasing, or insignificant Spearman correlation between summed expression and growth rate (Methods). Because noise leading to insignificant correlations can hide trends in summed expression, we use the following definitions for the R, C, and S sectors. The R sector included all clusters correlated positively with growth rate in at least one of the three sets of conditions (NaCl, TMP, or sugars) and not anti-correlated with growth rate in any condition. This included the classic ribosomal R sector genes, as well as all other non-ribosomal genes that rise with growth rate ^49,50^. The S sector included clusters anticorrelated with growth rate in NaCl or TMP but not in poor carbon (784 genes). The remaining clusters were defined as the C sector (1053 genes), which included clusters increasing in only poor carbon or in both poor carbon and NaCl. These sectors show expected overlap with previously reported sectors measured using proteomics^31^ (Fig. S12).

Enriched GO terms in these clusters (Fig. 3C, Methods) match the expectation that C is catabolism-related, R is ribosomal and growth-related, and S genes are stress-related. The inferred S sector also included anabolic genes, presumably related to requirements for production of protective components. For example, arginine biosynthesis is a known requirement for pH tolerance^51^. Because of their known role in antibiotic tolerance^52^, we measured the contribution of efflux pump expression to the observed protection from death. We found that the total expression of efflux pumps is very low and does not follow the expected expression trends of the model (Fig S13).

In accordance with the model assumptions, we fit the RNA-Seq sector-size data for sugars, NaCl, and TMP to linear functions of growth rate, requiring zero slope where sectors are expected to remain constant (S on sugars and C on stressors). We found an excellent fit in all three cases (Figs. 3D-F, Table S2; adj. R^2^=0.999 for sugars, R^2^=0.998 for NaCl, and R^2^=0.999 for TMP). Note that because sectors were defined using Spearman correlation, the linear dependence is not an artifact of the definitions. Notably, the R-sector slope is higher for sugars than for stressors. This difference may be due to a lower translation rate for R-sector genes under stress^53^, so that more ribosomal mRNA is required to provide a consistent amount of ribosomal protein. The ‘classic’ ribosomal R sector, which relates strictly to ribosomal content, can be measured by total RNA, which is predominantly ribosomal RNA. Total RNA shows the expected growth law: a linear dependence on growth rate that is similar in poor carbon sources and in stressors (Figs. 3F inset, S14, adj. R^2^=0.93)

### Experimental modulation of the C sector provides a rigorous test of the model

In Fig.2 we modulated the C sector by changing the carbon source. Another way to change the C sector and to further test the model is by modulating the activity of the master C-sector regulator CRP ^54^. This can be done by changing the concentration of the signaling molecule cAMP that activates CRP^32,35,36^. We thus measured growth and death rates of strain U486 (MG1655ΔcyaAΔcpdA), which cannot produce or degrade cAMP, in a range of exogenously supplied cAMP concentrations^32,35,36^.

As was shown earlier ^32,36^, growth rate is non-monotonic as a function of cAMP, yielding equi-growth rate conditions achieved by different cAMP levels (Fig S15). The optimum growth rate lies between 0.2-0.3mM cAMP. Death rate measurements yielded dependency on growth that did not collapse onto a single line, with low cAMP protecting more strongly from death compared to high cAMP at similar growth rates (Figs. 4A, S16, S17). Thus, as for the sugars and stressors, death rate as a function of cAMP is not determined solely by growth rate. Encouraged by the linear fits for sugars and stressors, we assumed that S remains constant for high cAMP (>∼0.3mM cAMP) while C remains constant for low cAMP (<∼ 0.2mM cAMP). Strikingly, despite the nonlinear growth curve as a function of cAMP, this same model provides an excellent fit (adj. R^2^=0.941) with just two slope parameters (Table S3).

**Fig. 4.**
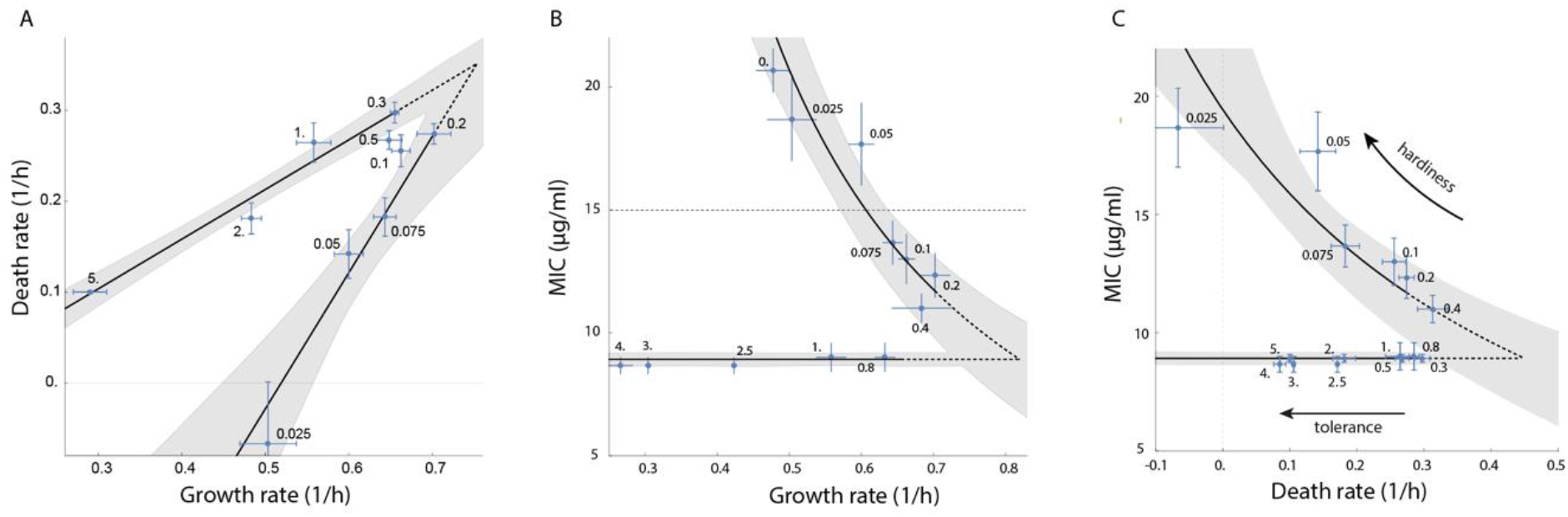
The proposed resource allocation model quantitatively captures death rate and MIC as a function of cAMP in a ΔcyaA ΔcpdA strain. (**A)** Death rate as a function of growth rate upon treatment with 15μg/ml nalidixic acid fit with two slope parameters. (**B**) MIC as a function of growth rate fit with one anchor parameter. Labels are cAMP concentration in mM. Shaded regions represent 90% confidence intervals on the fit. Dashed portions of fits are extrapolations into the unmodeled transition region between low and high cAMP regimes. Dashed fit lines represent extrapolations into the unmodeled transition region between low- and high-cAMP regimes. Horizontal dashed line in **B** is the concentration of nalidixic acid used in **A. (C)** Plotting MIC vs death rate for emphasizes the difference between tolerance and hardiness. High cAMP concentrations show tolerance (decreased death rate without a change in MIC), while low cAMP concentrations show hardiness (decreased death with a corresponding increase in MIC). For data in which only MIC or death rate was measured in **A** and **B**, the other was imputed from the fit. Note that the maximum growth rate in this strain differs from the wild type used in Figs 2-3.

We also tested the MIC in these conditions, and again find different relationships between MIC and growth rate in the low and high regimes of cAMP levels (Fig. 4B). For cAMP>∼0.3mM MIC was constant with growth rate, while for cAMP <∼0.2mM, it increased with decreased growth rate. Using one additional free parameter to describe the anchor point between the two regimes fit the data very well (adj. R^2^=0.991, Methods). Plotting MIC versus death rate highlights the difference between tolerance and hardiness: high cAMP shows tolerance while low cAMP shows hardiness (Fig. 4C).

The origin of the differential behavior in low and high cAMP regimes is revealing. The present model indicates that at cAMP levels above the optimal growth rate, the C sector rises with cAMP ^32,36^ at the expense of the R sector without a change in the S sector, similar to the sugar line. Death drops with cAMP, as does growth. At cAMP levels below the optimal growth rate, the C sector changes only mildly ^32^, and the S sector rises with decreasing cAMP at the expense of the R sector, with a protective effect on death. Thus, ultra-low cAMP levels, achieved physiologically only in unusual conditions^55^, are interpreted by the cells as a stress signal, leading to a reduced death rate and an increased MIC.

## Discussion

We developed a high-throughput assay for measuring bacterial death curves. Using this assay, we determined the relation between the growth rate in a given condition and the death rate in a subsequent antibiotic challenge. Death rate depended both on growth rate and growth condition. Stressful conditions protected from death when compared to no-stress conditions of equal growth rate. Stress resulted in lower death rate and increased MIC, a phenomenon that lies outside the available definitions of antibiotic response, which we termed hardiness. The quantitative relation between growth and death is captured by a resource allocation model, in which death is increased by growth-related damage and reduced by a protective stress-response sector. We identified this stress sector using RNA-Seq measurements. In line with the model, protection from death was also gained when manipulating the C-sector to low levels via exogenous cAMP.

Most of the work on bacterial growth laws has focused on growth rate, not on death rate. Growth laws are explained by resource allocation models that focus on the relation between proteome distribution and growth rate. These models define sectors by an increased expression of a group of genes needed to cope with a certain limitation at the expense of ribosomes ^30–32^. Here, we find that resource allocation-based theory can also predict death rate when we introduce a newly defined stress-response sector.

These findings emphasize the need to define what conditions are considered stressful for bacteria. Stress was recently defined as a condition that limits growth not by nutrients but through other environmental parameters^34^. Such conditions will upregulate a proteomic response that is not directly involved in biomass production (such as ribosomes, catabolic genes). One of the stressors we used is the bacteriostatic antibiotic TMP. Our results thus provide an explanation for the known antagonistic effect of bacteriostatic antibiotics on bactericidal action^56^: bacteriostatic antibiotics in general may raise a stress response and thereby reduce death by bactericidal antibiotics. An additional way to impose stress is starvation, known to provide tolerance and persistence to antibiotics while upregulating stress-gene expression^9,10,23,57^. It will be interesting to measure antibiotic-mediated death rates and the size of the S sector under starvation. Likewise, it will be interesting to measure the effect of nitrogen limitation on various carbon sources, given that various combinations of carbon and nitrogen sources can lead to either low or high cAMP levels ^55^. It may also be illuminating to explore the connection of the proposed coarse-grained model of stress to molecular regulators of stress response such as ppGpp^58^.

In addition to the connection between growth conditions and death rate, the present findings highlight the importance of considering growth conditions when determining the MIC for a given antibiotic. An elevated MIC is usually considered a form of resistance, such as that caused by mutations. A raised MIC can also result from stressful growth conditions without mutations^23,59^, in a manner captured quantitatively by the present resource allocation model. Our proposed differentiation between tolerance and hardiness captures the range of behaviors that describe both MIC and death rate.

Clinically, our work suggests that in the variety of conditions in the body, bacteria may be tolerant and hardy to antibiotics compared to laboratory conditions. For example, some of the stress placed on bacteria by the immune system may counterintuitively inhibit the killing efficacy of antibiotics. This suggests possible targets for treatments. For instance, provision of inhibitors of alternative sigma factors together with antibiotics may inhibit stress sector expression and enhance antibiotic efficacy. We anticipate that quantitative understanding of the death-growth tradeoff in bacteria and its relation to stress may thus have clinical applications, and more generally may advance our understanding of tradeoffs in bacterial physiology.

## Methods

### Strains

Experiments in this study were carried out using either NCM3722 strain (CGSC #12355) or MG1655 strain (CGSC #6300) and its derivatives MG1655ΔcyaA/ΔcpdA (U486^36^). All strains were transformed with a kanamycin resistant, low copy promoterless plasmid (U66^60^) to acquire resistance and reduce contaminations during the long death experiments.

### Growth rate and MIC measurements

Cells were grown overnight in M9 minimal medium (42mM Na_2_HPO_4_, 22mM KH_2_PO_4_, 8.5mM NaCl, 18.5mM NH_4_Cl, 2mM MgSO_4_, 0.1mM CaCl) containing 11mM glucose, 0.05% casamino acids and 50μg/ml kanamycin at 37°C and diluted 1:300 into the designated Media (all based on M9 with different carbon source, different concentration of NaCl etc.)

For growth rate measurements cultures were distributed using a robotic liquid handler (FreedomEvo, Tecan) in 96-well plates (150μl per well in at least 6 wells). Wells were covered with 100 µl of mineral oil (Sigma) to prevent evaporation and transferred into an automated incubator. Cells were grown in an automated incubator with shaking (6 hz) at 37°C for about 20 hours. Every 6-10 minutes the plate was transferred by a robotic arm into a multi-well fluorimeter (Infinite M200Pro, Tecan) that reads the bacteria optical density (OD, 600nm). For MIC measurements the setup was very similar with the following changes: cultures were pre-grown in tubes for 3-4 hrs and then distributed to wells containing different concentrations of Nalidixic acid.

Growth rate was calculated as the temporal derivative of the natural logarithm of the OD curves, *µ* = *dln*(*OD*)/*dt*. Exponential growth rate is the mean over a region of at least 2 generations with a nearly constant growth rate.

MIC was determined as the minimal Nalidixic acid concentration which led to an OD decline (in NCM3722 strain, Fig. S6A) or a non-increasing OD curve (MG1655 strain, Fig. S6B).

### Death rate measurements

A scheme of the experimental setup (carried out automatically in a robotic system (FreedomEvo, Tecan)) is shown in Fig 1 for one growth condition and a short experiment duration. In reality, in each experiment we measured 6-8 growth conditions and up to 16 hrs of antibiotic challenge.

This experimental setup contains 3 stages:

1. *Pre-growth stage*: This stage was carried out in a 96 well plate containing 200μl media (3-4 different growth conditions in a plate) +50μl mineral oil. As shown in Fig.1 an overnight culture (M9+11mM glucose+0.05% casamino acids+50μg/ml kanamycin) was diluted (1:300) into the growth plate. The culture was diluted to the first well of each condition (wells A1, A4, A7, A10 for a 4-conditions plate) and incubated in an automated incubator with shaking (6 hz) at 37°C for a time period (0.5-1 hr) which defines the experiment time resolution. Plates were transferred by the robotic arm into a multi-well fluorimeter (Infinite M200Pro, Tecan) that reads OD (600nm), followed by bacteria transfer from the overnight culture to the successive wells. This stage was repeated till the first well reached exponential phase (3-8 hrs, OD 0.02-0.04).
2. *Antibiotic-challenge stage*: This stage was carried out in a 96 well plate containing 170μl media (same growth conditions as in the pre-growth plate+ Nalidixic acid) +50μl mineral oil). The first well of each condition of the pre-growth plate is diluted (1:7) into the antibiotic-challenged plate, in parallel inoculation of the ON culture to the pre-growth plate continues as well. Both plates are incubated with shaking (6 hz) at 37°C for the same time used in the pre-growth stage. Plates were transferred into a multi-well fluorimeter for OD measurements, followed by bacteria transfer to successive wells. For experiments with time resolution of 1 hr this stage was routinely repeated 16 times. For experiments with time resolution of 0.5 hr more repeats were carried out.
3. *Recovery stage*: In the last stage we adopted the protocol of Hazan *et. al*.^40^ for viable cell determination based on the incubation time to cross a certain OD threshold. After the last transfer from the pre-growth plate to the challenged plate we immediately transferred the challenged plate to ice. Bacteria treated for different times as well as a non-treated culture were diluted 1:100 into 1 ml of recovery medium (M9+11mM glucose+50μg/ml kanamycin) in a deep 96-well plate. Non-treated cultures were also serially diluted in order to obtain a standard curve. Using the robotic system, we transferred each diluted culture to 6 wells of a 96-well plate (150μl per well). Wells were covered with 100 µl of mineral oil and transferred into an automated incubator. Cells were grown in an automated incubator with shaking (6 hz) at 37°C for about 20 hours. Every 10 minutes the plate was transferred by a robotic arm into a multi-well fluorimeter (Infinite M200Pro, Tecan) that reads the bacteria optical density (OD, 600nm). Setting the OD threshold to 0.05, (∼2 · *background OD*), we extracted from each OD curve the time (*τ*) required to reach this threshold (Fig1. B). Using the standard curve (Fig. 1C) we obtained for each growth condition the fraction of surviving bacteria as a function of antibiotic treatment time (Fig. 1D).
4. *Death rate calculation*: For each condition, we obtained an average survival curve based on at least 3 biological repeats (for the cAMP data we measured more cAMP levels with less repeats on each level). We fit the data with a Weibull survival function coupled to exponential growth, 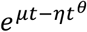, which allows fitting of exponential and sigmoidal survival curves as well as allowing for an initial increase in viable cells. For conditions in which the maximum occurred at 0 hr, we assumed *µ* = 0 to ease fitting. To further ease the nonlinear fit, we restricted *µ* < 1 (since growth rate without antibiotics are all below 1), *η* > 1 (at least exponential decay), that the maximum is below 2 and the time to reach the maximum is less than 2 hrs (well consistent with all data). We define death rate as 1/time for the fit to lose 90% of the population compared to the initial value at 0 hr. For cultures that grew, we defined a negative death rate as -1/time for the fit to reach 10x the initial values at 0 hr. Values were averaged over the number of biological replicates indicated in Figs. S2 and S16, with errors given as the standard error of the mean. R^2^ values for all fits across all biological replicates are provided as a histogram in Fig. S18. For presentation purposes, additional average survival curves are shown in Figs. S3 and S17, with additional Weibull plus growth fits. The R^2^ values for these fits are provided as a histogram in Fig. S19.

### RNA sequencing

Cultures for RNA-seq were grown to exponential phase to OD lower than 0.1 in microplates, to match the OD at which antibiotic was added to the growing cultures in the death assay. RNA was extracted from these exponentially growing cultures using RNAeasy Protect bacteria Mini Kit (Qiagen). Total RNA was measured using Nanodrop (Thermo Scientific). For RNA sequencing rRNA was depleted using NEBNext rRNA depletion kit. RNAseq libraries were prepared at the Crown Genomics institute of the Nancy and Stephen Grand Israel National Center for Personalized Medicine, Weizmann Institute of Science. Libraries were prepared using the INCPM-mRNA-seq without polyA selection protocol. Briefly, 80 ng of input RNA after ribosomal depletion was used for fragmentation and generation of double-stranded cDNA. After Agencourt Ampure XP beads cleanup (Beckman Coulter), end repair, A base addition, adapter ligation and PCR amplification steps were performed. Libraries were quantified by Qubit (Thermo fisher scientific) and TapeStation (Agilent). Sequencing was done on a Nextseq instrument (Illumina) using a 75 cycles high output kit. The Reads were mapped to the MG1655 genome. Expression levels for each gene were quantified using htseq-count.

### RNA-Seq analysis and clustering

We acquired data for 4229 genes across all measured conditions, accounting for ∼97% of *E. coli* genes. Because we sum gene expression to find sector sizes, we did not filter out low-read count genes. All raw count data was normalized using DeSeq2 after filtering out residual rRNA and tRNA reads. The Mclust ^61,62^ library was used to perform a Gaussian mixture model. Mclust recommended between 10 and 30 clusters with similar Bayesian Information Criterion (BIC) and spherical, unequal volume model (“VII”). We used the VII model with 12 clusters. The Spearman correlation for sugars, NaCl, and TMP conditions was subsequently calculated within each cluster, categorizing each line in each cluster as correlated, anticorrelated, or unchanging with growth rate. A liberal p-value cutoff of 0.2 for the correlation was used so as to include all genes as changing significantly in at least one set of conditions in at least one cluster. Clusters with only unchanging or positive correlations with growth rate were categorized as R. Clusters not in R and not anticorrelated with growth rate of the carbon sources were categorized as S. The remaining clusters were categorized as C, which included to some extent clusters increasing both in poor carbon sources and in increasing NaCl concentration.

### Gene Ontology Analysis

Gene lists for each of the three sectors were checked for biological process gene ontology significance against the full list of E. coli genes using the rbioapi interface for PANTHER ^63^ using default parameters, including a significance cutoff of false discovery rate (FDR) below 0.05. The full list of GO terms and their significance are provided in Dataset S1. The terms displayed in Fig. 3D are those 5 terms with no significant encompassing terms with the lowest FDR.

### Nondimensionalization

The underlying equations *R* + *C* + *S* = 1, *µ* = *aR* − *b*, and *ρ* = *αµ* − *βS* + *ϵ* were converted to glucose-relative Eqs 1-3 in the main text by subtracting off equations *R*_*G*_ + *C*_*G*_ + *S*_*G*_ = 1, *µ*_*G*_ = *aR*_*G*_ − *b*, and *ρ*_*G*_ = *αµ*_*G*_ − *βS*_*G*_ + *ϵ* where the G subscript indicates glucose. The deltas in the main text are defined as, e.g., Δ*µ* = *µ* − *µ*_*G*_. For fitting, a further simplification was made by non-dimensionalizing. Specifically, we define 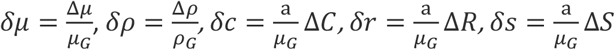.

This yields non-dimensionalized equations

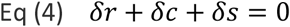

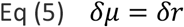

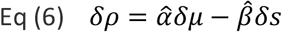

where 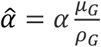 and 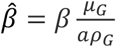.

### Curve fitting

All fits in the paper were performed in Mathematica using LinearModelFit or NonlinearModelFit.

For Fig. 3A, Eqs 4-6 were solved for death rate as a function of growth rate at either *δc* = 0 (stressors) or *δs* = 0 (sugars), eliminated for *δr*. This yielded the following prediction for death rate:

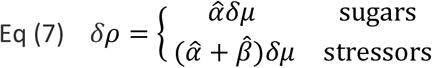

Data was organized as 3-value points (type index, growth rate, death rate), with the index specifying the data point as sugar, NaCl, or TMP. These data were non-dimensionalized using the measured values for glucose and fit to Eq. 7. Fitting weights were given as the inverse of the sum-square error of growth and death rates, with growth and death errors normalized first across all samples. Glucose was included as both a sugar, NaCl, and TMP, with 1/3-weight each. Because we assume measured values for the growth and death rates on glucose, there are two sources of error on the fit. The first derives from the fitting error when glucose values are given and the second derives from the variation in the glucose measurements themselves. We thus calculate error bands for fits as 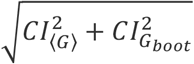. Here *CI*_⟨*G*⟩_ is defined as the 90% confidence interval of the fit with glucose specified as the average glucose measurements. 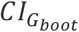 is defined as the 90% confidence interval across 1000 fits, where for each fit the glucose growth and death rates were sampled from a Normal distribution with mean and standard deviation given by the measured average and standard error of glucose displayed in Fig. 2. The same procedure was followed in Figs. 3D-F, 4A, and 4B (low cAMP). Error bands for Figs. 3B and 4B (high cAMP) were produced using the same procedure where already fit parameters were likewise included in the bootstrapping.

For Fig. 3B, we assumed that growth-derived death *α* = *α*_0_(*m* − *m*_*G*_) and glucose death rate *ρ*_*G*_ = *ρ*_0_(*m* − *m*_*G*_) increased linearly with antibiotic concentration *m*, relative to a reference *m*_*G*_. Given that *m* equals the MIC when *ρ* = 0 for any condition, we immediately see that *m*_*G*_ is the MIC on glucose. Substituting these relations into Eq. 7 yields the following prediction for the MIC (*m*) for all conditions when setting *ρ* = 0:

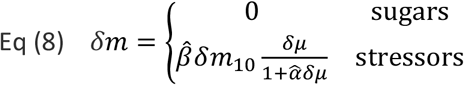

with 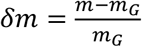 and 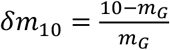 comes from the fact that 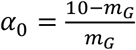, the value 10 being the antibiotic concentration used in fitting the values of alpha and beta in Fig. 3A. Note that the constant value in sugars is not assumed upfront, but rather derives from the assumption that Δ*S* = 0 in sugars.

For Fig. 3E, a separate fit was performed for sugars, NaCl, and TMP. In each case, the data were fit to the expected piecewise linear function implied by Eq. 2:

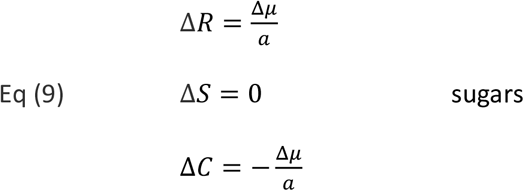

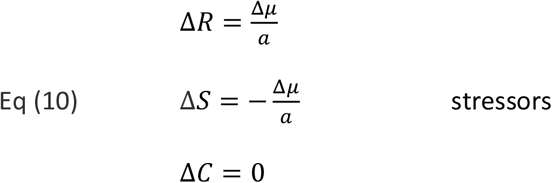

These equations have three free parameters, *C*_*G*_, *S*_*G*_, and *a*, where the glucose growth rate reference is taken as given from the data as described above for the fit in Fig. 3A.

For the relationship between total RNA and growth rate, we fit

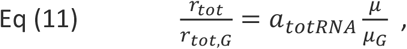

where *r*_*tot*_ is the total RNA, *r*_*tot,G*_ the total RNA in glucose growth, *µ* the growth rate, *µ*_*G*_ the growth rate on glucose, and *a*_*totRNA*_ the fitted slope. We performed the fit for sugar, NaCl, and TMP conditions separately, as well as with all data combined.

For Fig. 4A, the same procedure was used as in Fig. 3A, but a separate anchor was used for low cAMP data (0.2mM cAMP) and high cAMP data (0.3mM cAMP), as described in the main text, with new parameters 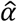 and 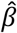:

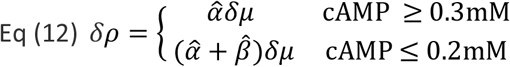

For Fig. 4B, *m*_*G*_ was estimated directly as the average of MIC values for measured points at cAMP > 0.4mM. A separate, extrapolated anchor point (“pseudoglucose” growth rate *µ*_*PG*_) was fit for to the low cAMP data using Eq 8, with *δm*_10_ replaced with 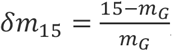, the value 15 being the concentration of nalidixic acid used in Fig. 4A:

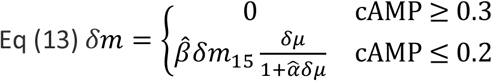

## Code availability

RNA-Seq analysis was performed in R version 4.1.0. All other fitting and analytical manipulation were performed in Mathematica version 13. Source code can be found in Dataset S2.

## Data availability

All data not provided in the text or supplements is available upon request.

## Acknowledgements

We thank B. Towbin, S. Kostinski, T. Milo, A. Bar, Y. Yang, and E. Vaisbourd for comments on the manuscript. Funding was provided by European Research Council (ERC) under the European Union’s Horizon 2020 research and innovation program (grant agreement No. 856587). D.S.G. was funded as a member of the Zuckerman Postdoctoral Scholars Program. U.A is the incumbent of the Abisch-Frenkel Professional Chair.

## Author contributions

A.B. conducted experiments and analysis. A.M. and D.S.G. performed theoretical analysis and fits. All authors contributed to conception and to writing of the manuscript.

## Supplementary information

Supplementary Figs. S1-S11 and Tables S1-S2 are provided in the supplementary information.

