## Supplementary Information for "Tradeoffs in bacterial physiology determine the efficiency of antibiotic killing"

Anat Bren*, David S. Glass*, Yael Korem Kohanim, Avi Mayo, Uri Alon
Department of Molecular Cell Biology, Weizmann Institute of Science, Rehovot, Israel

**Index:**

**Figure S1.** Cultures were exposed to antibiotics during exponential growth at matching ODs 2
**Figure S2.** All survival curves used in this study for the NCM3722 strain as obtained by our automatic death assay 3
**Figure S3.** Weibull death plus growth for averaged survival curves in NCM3722 for presentation purposes 4
**Figure S4.** Death curves obtained by our assay agree well with curves obtained by the CFU method 5
**Figure S5.** Calibration curves of dilution vs. recovery lag time do not depend on time in antibiotic 6
**Figure S6.** Survival fraction of day-to-day repeats of cultures grown under high-stress conditions 7
**Figure S7.** Stressors with growth rates near or below that of mannose show significantly lower death rate 8
**Figure S8.** Stressors NaCl and TMP protect against death in phosphomycin and streptomycin 9
**Figure S9.** Conditions expected to supply both stress and poor carbon lie between sugar and stress curves 10
**Figure S10.** Growth curves are used to determine the MIC 11
**Figure S11.** MIC vs death rate for NCM3722 data emphasizes the difference between tolerance and hardiness 12
**Figure S12.** Comparison of RNA-Seq data from this work to proteomic measurements in previous work 13
**Figure S13.** Efflux pump expression is a small fraction of the S sector 14
**Figure S14.** Total RNA as in Fig 3F inset without normalization to glucose 15
**Figure S15.** Growth rate as a function of cAMP in MG1655ΔcyaAΔcpdA is a non-monotonic function 16
**Figure S16.** All survival curves for MG1655 ΔcyaA ΔcpdA (U486) strain as obtained by our automatic death assay 17
**Figure S17.** Weibull death plus growth for averaged survival curves in U486 for presentation purposes 18
**Figure S18.** $R^2$ for Weibull plus growth fits for all NCM3722 and 486 survival curves in Naladixic acid 19
**Figure S19.** $R^2$ for Weibull plus growth fits for all NCM3722 and U486 averaged survival curves in Naladixic acid 20
**Table S1.** Parameter values for Fig. 3A 21
**Table S2.** Parameter values for Fig. 3D-F 21
**Table S3.** Parameter values for Fig. 4A 21
**Table S4.** Parameter values for Fig. 4B 21
**Supplemental references** 21

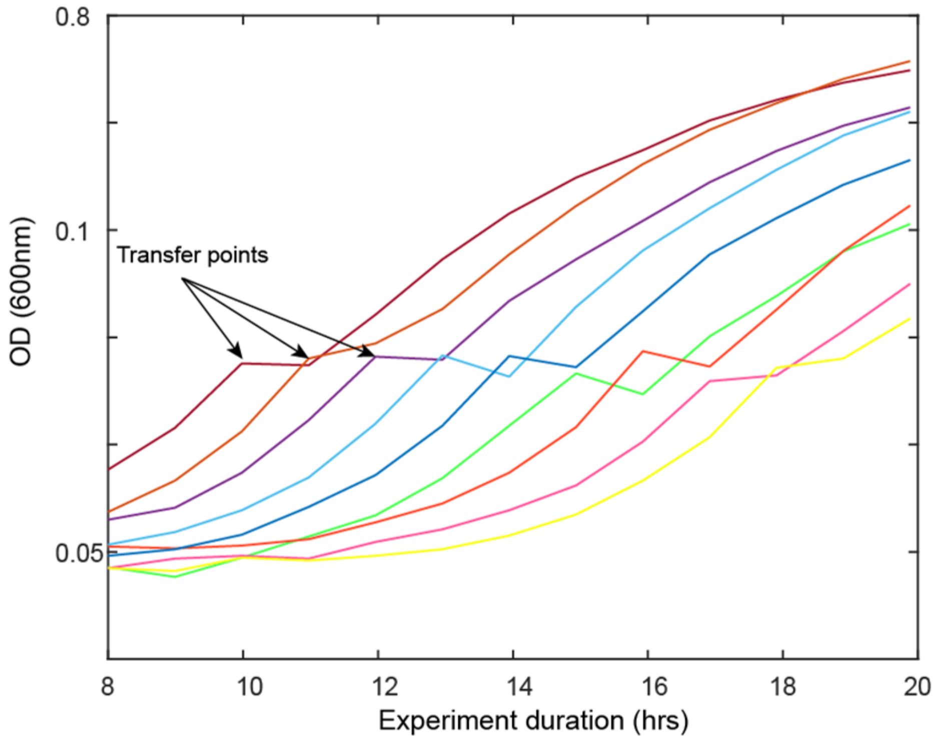


**Figure S1. Cultures were exposed to antibiotics at similar ODs during exponential growth.** Each curve represents the OD of wells inoculated at successive times in the pre-growth plate. The shift of each curve is due to the change in volume of the growth-well contents when a sample was transferred to the challenge plate, and thus indicates the time at which the culture was transferred to the antibiotic challenge. This transition takes place at exponential phase and at a similar OD for all wells, indicating that a similar number of bacteria at the same growth stage were challenged by the antibiotics throughout the experiment.

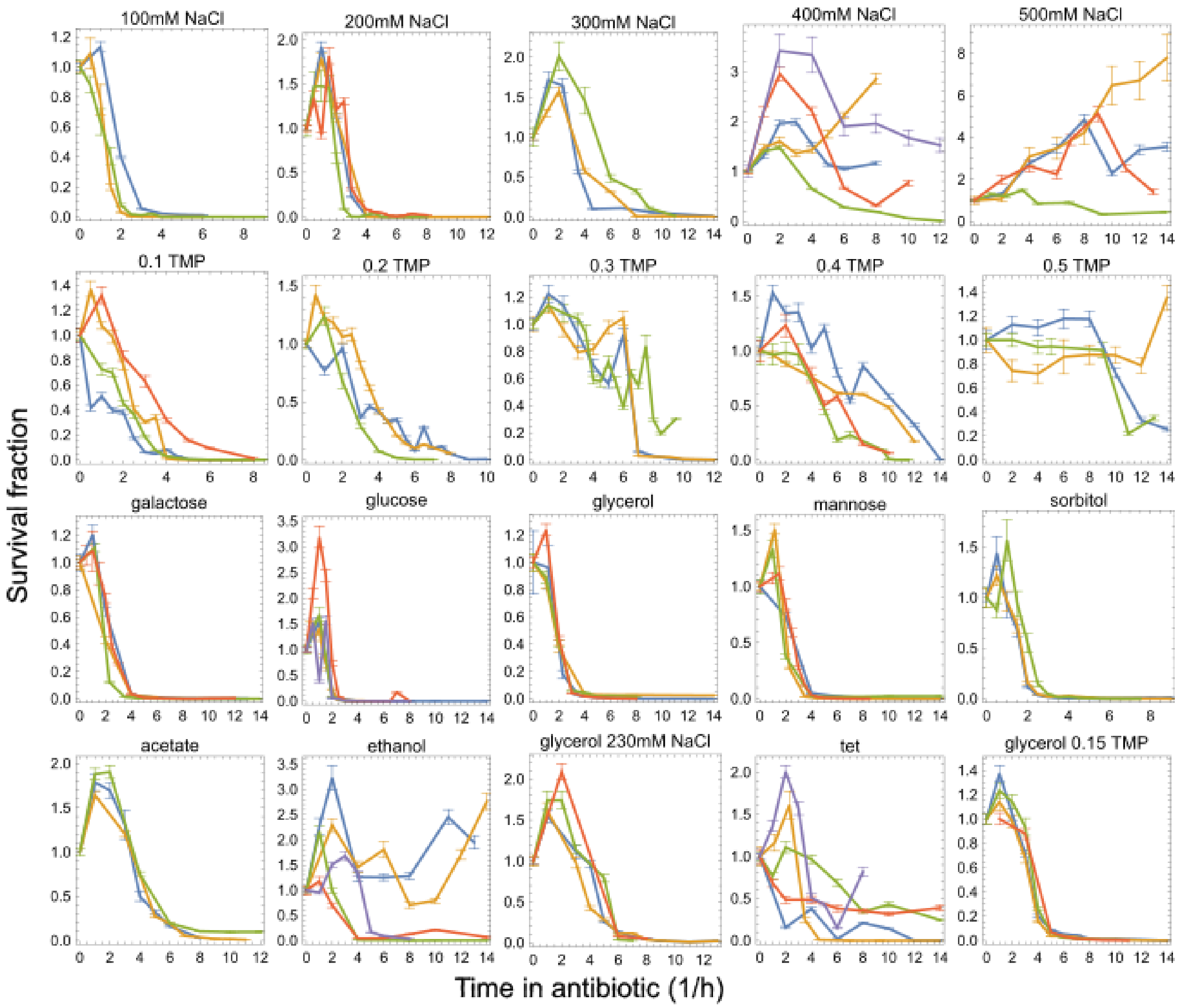


**Figure S2. All survival curves used in this study for the NCM3722 strain as obtained by our automatic death assay.** Cultures were challenged with 10µg/ml Nalidixic acid. Different colors in each panel are biological repeats, almost always performed on separate days.

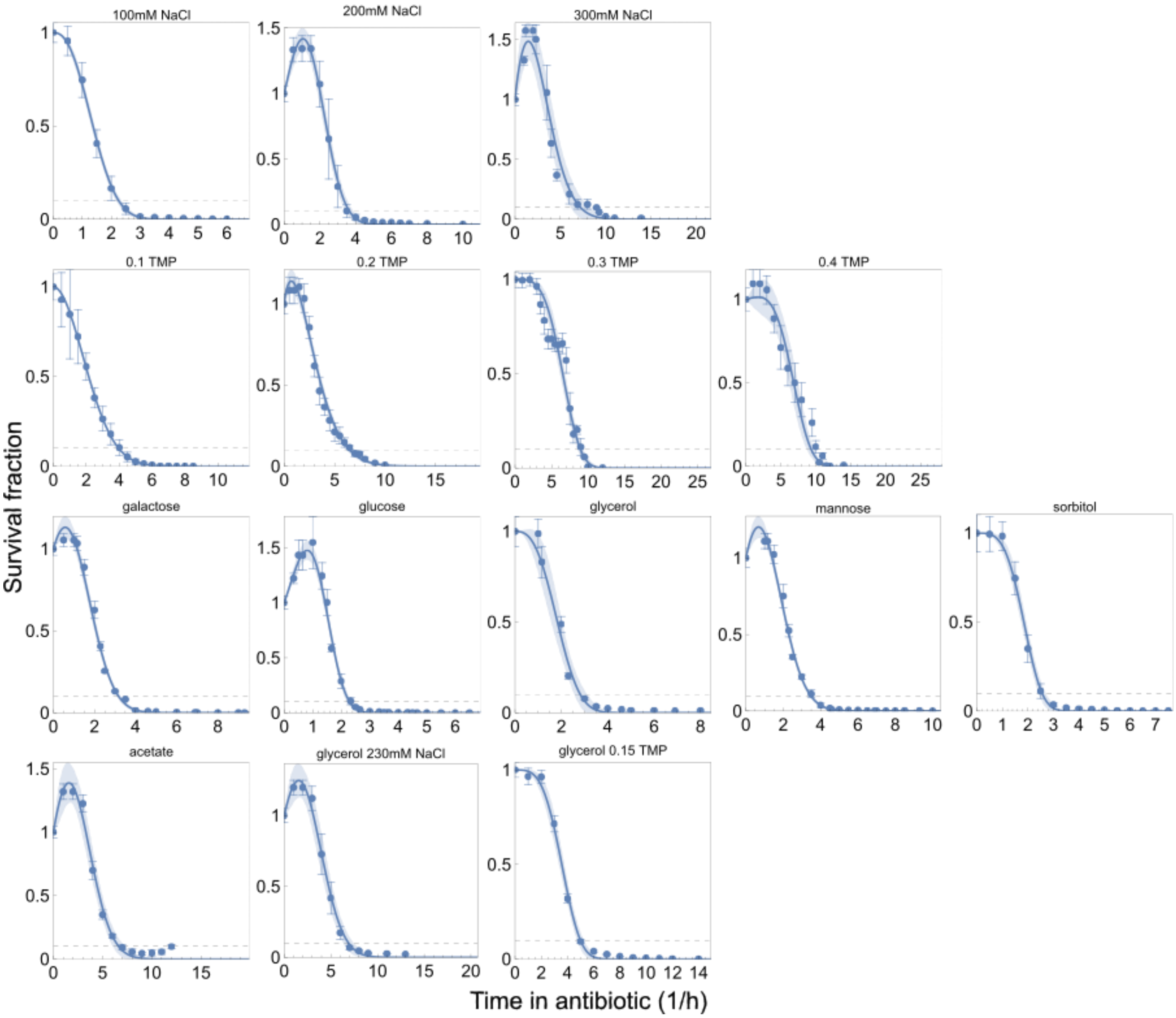


**Figure S3. Weibull death plus growth for averaged survival curves in NCM3722 for presentation purposes.** Dashed line represents 10% survival. Shaded bands represent 95% confidence intervals on the fit. Points here are weighted averages of the data in Fig. S4. Death rates were calculated from similar fits to the individual replicates in Fig. S4 and then averaged.

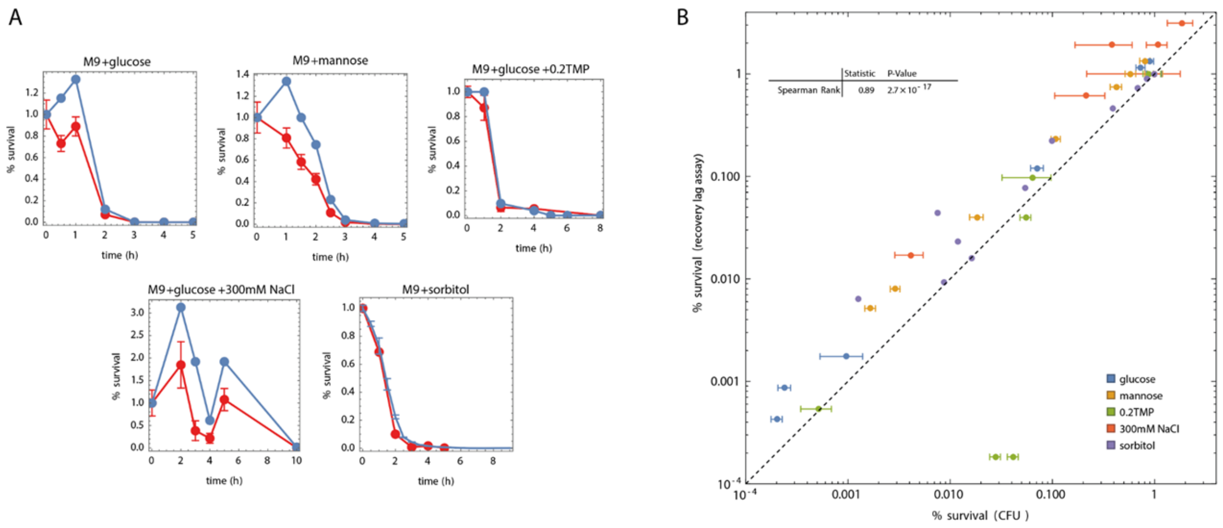


**Figure S4. Death curves obtained by our assay (blue line) agree well with curves obtained by the CFU method. (A)** The blue dots were obtained by the recovery lag time protocol described in Fig. 1, and the red dots were obtained by plating the same treated cultures on LB agar plates+50µg/ml kanamycin. Note that the initial increase seen in our assay is not recapitulated in CFU, likely indicating cell growth without division. This may correspond to known filamentation responses under antibiotic treatment[1] . **(B)** We found a high correlation (0.89) between all data points collected by recovery lag vs. CFU.

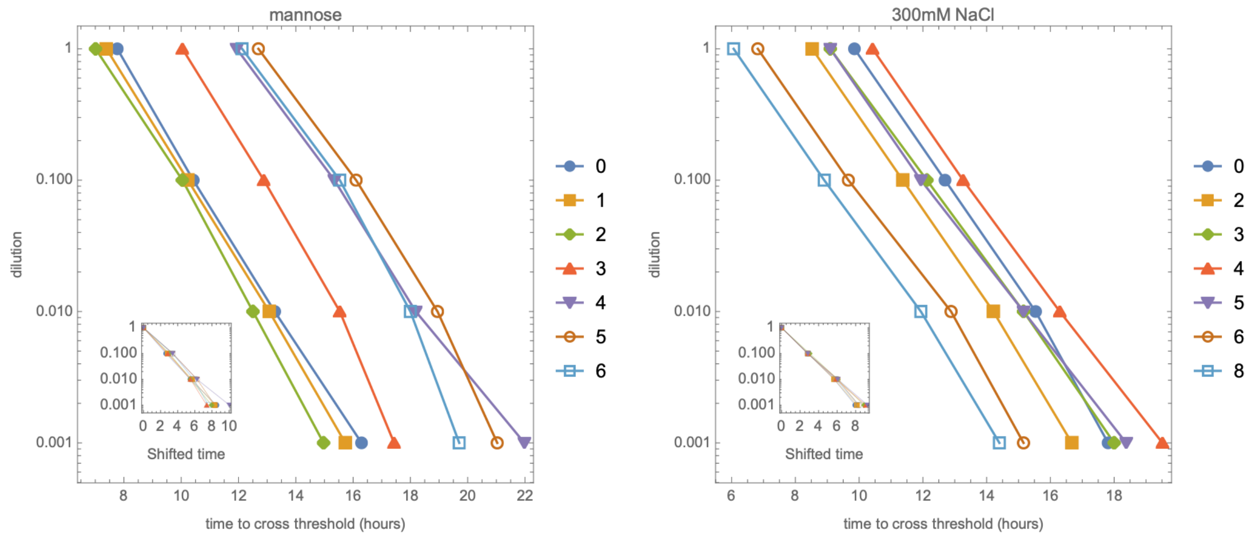


**Figure S5. Calibration curves of dilution vs. recovery lag time do not depend on time in antibiotic.** Increased time in antibiotic (shown by different colors) shifts the lag without dilution due to fewer surviving cells, but normalization shows that the slopes are invariant to time in antibiotic (insets).

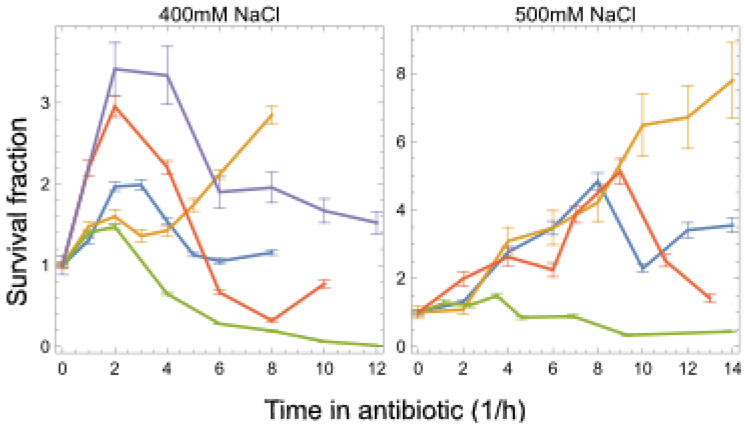


**Figure S6. Survival fraction of day-to-day repeats of cultures grown under high-stress conditions.** Stress conditions were 400mM and 500mM NaCl, Note that cells grew faster than they died in such conditions, leading to rising and noisy survival curves. Rising curves were assigned a negative death rate (see Methods).

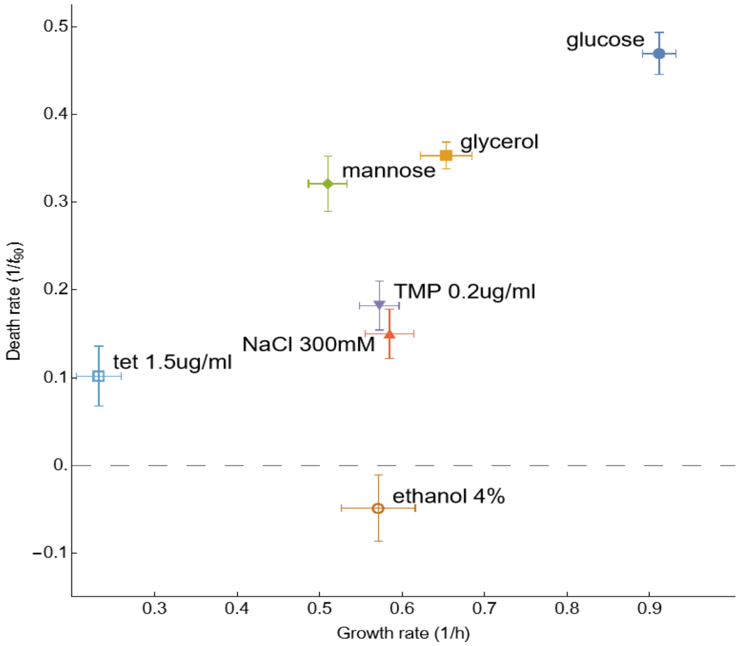


**Figure S7. Stressors with growth rates near or below that of mannose show significantly lower death rate.** Here we include the additional stressors ethanol and tetracycline, in addition to the NaCl and TMP shown in the main text.

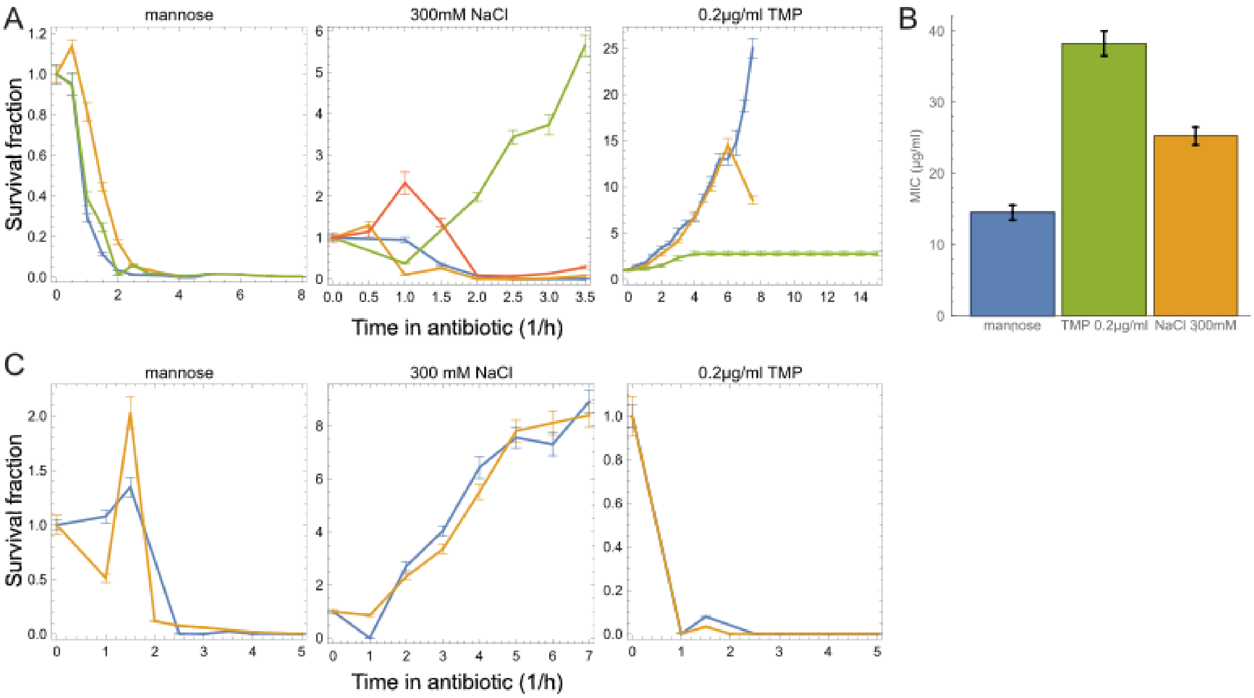


**Figure S8. Stressors NaCl and TMP protect against death in phosphomycin and streptomycin. (A)** Survival curves in 30µg/ml streptinomycin, showing that 300mM NaCl and 0.2µg/ml TMP have lower death rates than mannose. which have similar growth rates. **(B)** The MIC in 15µg/ml streptinomycin of the TMP and NaCl cultures was significantly higher than under mannose growth. **(C)** Survival curves in streptomycin showing that 300mM NaCl protects against death while 0.2µg/ml TMP does not.

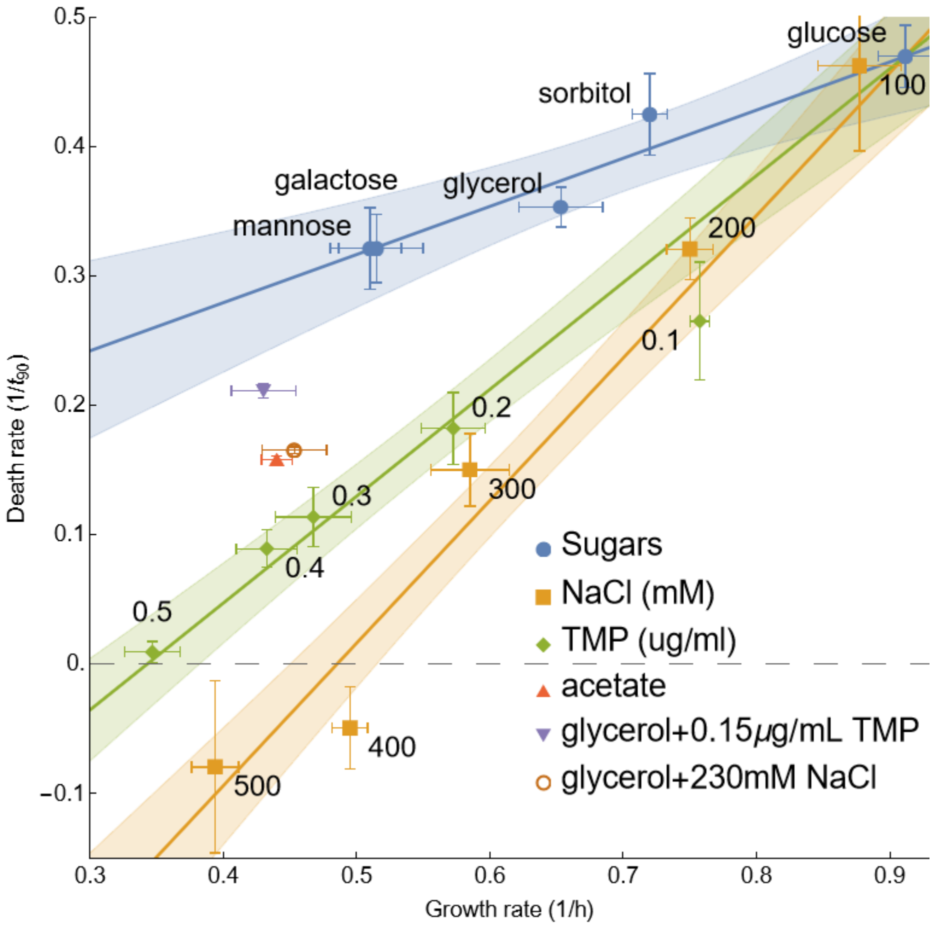


**Figure S9. Growth conditions expected to supply both stress and poor carbon lie between the sugar and stress curves, as expected.** Note that acetate is both a carbon source and a stressor.

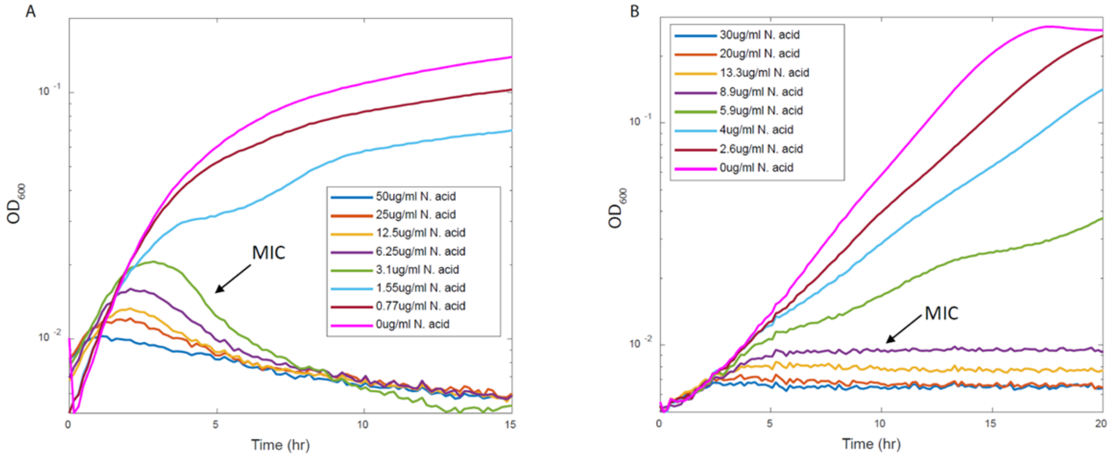


**Figure S10. Growth curves were used to determine the MIC.** Example plots in (**A**) NCM3722 and (**B**) MG1655 strains. Cells were grown under different conditions and a range of nalidixic acid concentrations. MIC was determined as approximately the concentration leading to a flattening growth curve.

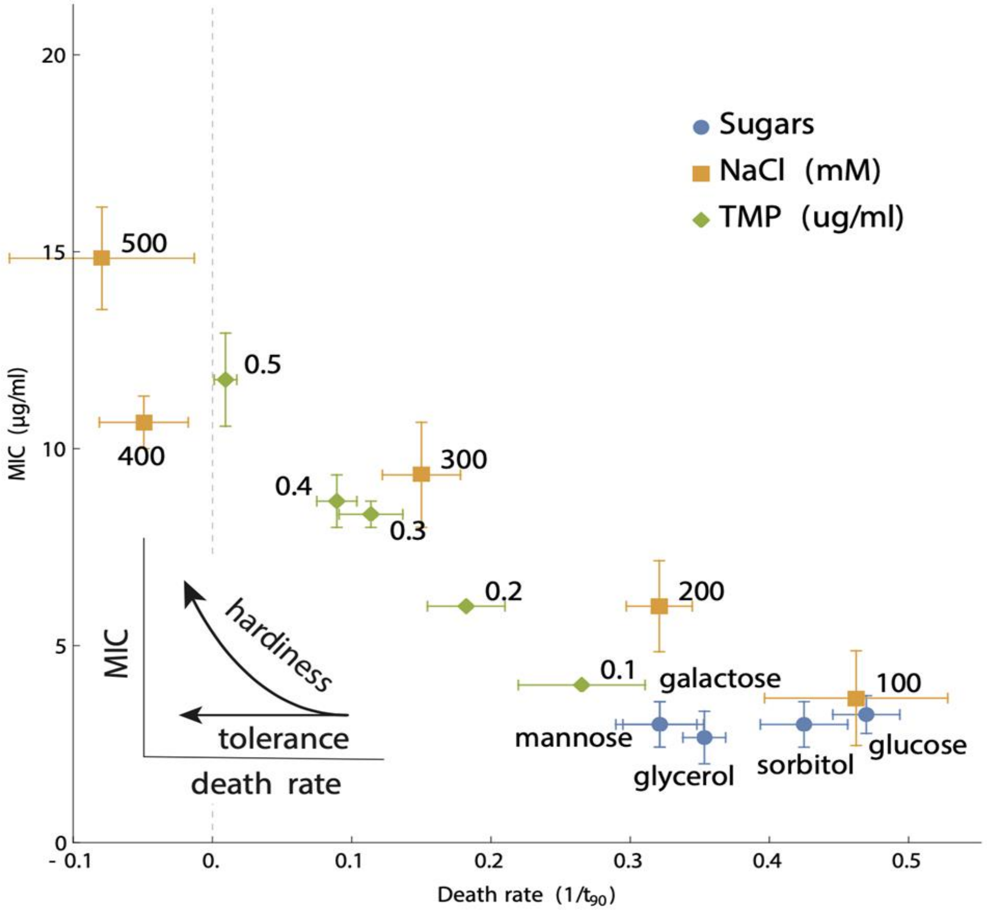


**Figure S11. MIC vs death rate for NCM3722 data emphasizes the difference between tolerance and hardiness.** Sugars show tolerance (decreased death rate without a change in MIC), whereas stressors show hardiness (decreased death with a corresponding increase in MIC).

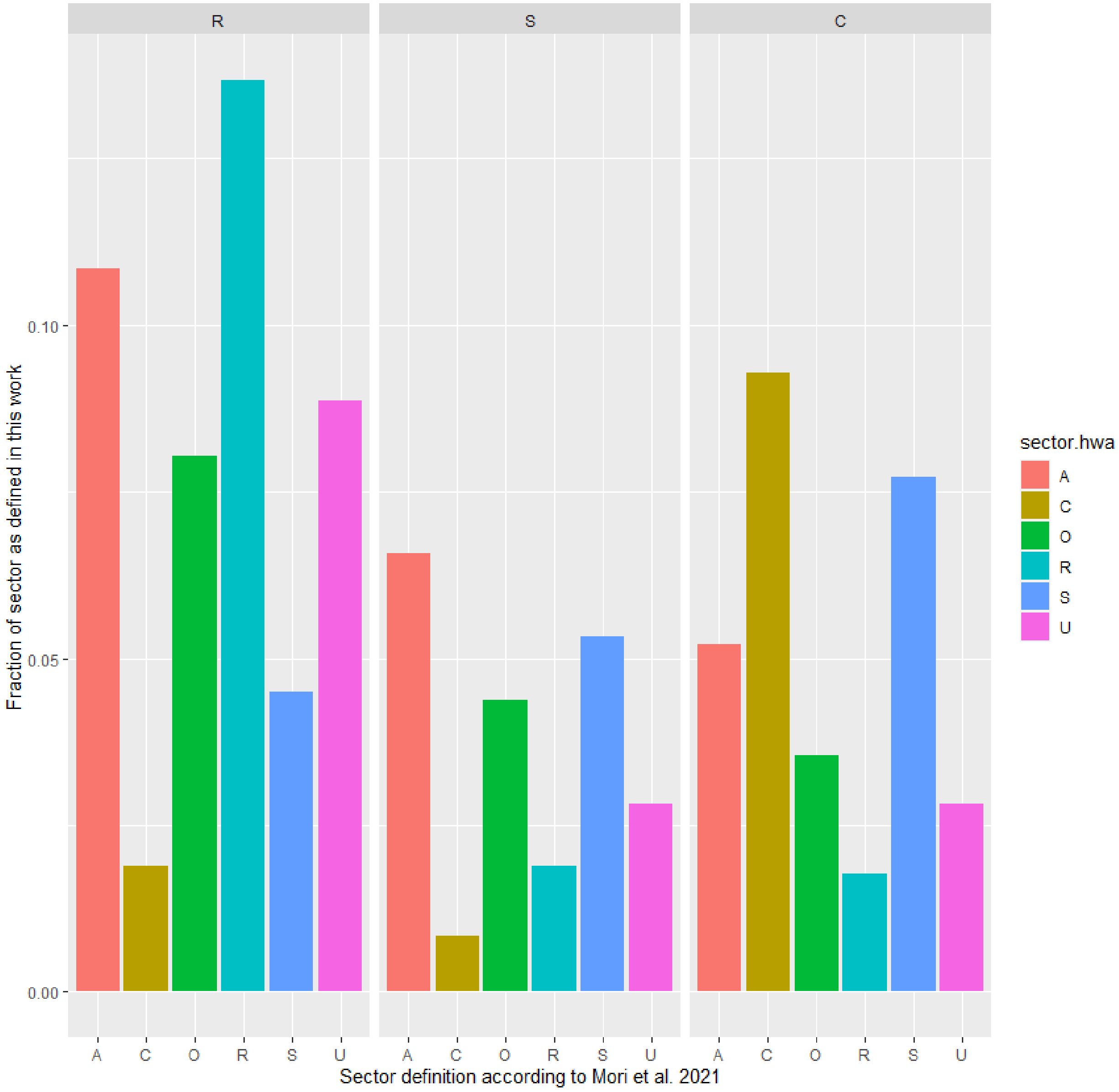


**Figure S12. Comparison of RNA-Seq data from this work to proteomic measurements in previous work.** In Hui *et al.*[2], R corresponds to ribosomal limitation (via chloramphenicol treatment), C increases under catabolic limitation, A under anabolic limitation, and S under both anabolic and catabolic limitation. U increased under all three limitations and O had more complex trends. Note that the "S" defined here is not equivalent to the "S" defined in Hui *et al*. We expect that the C sector in this work should overlap primarily with C and S and the R sector with R, A, and S. Because Hui *et al*. did not measure in stress conditions, we expect our S definition to be distributed more widely over their sectors. The comparison here shows qualitative support of that expectation.

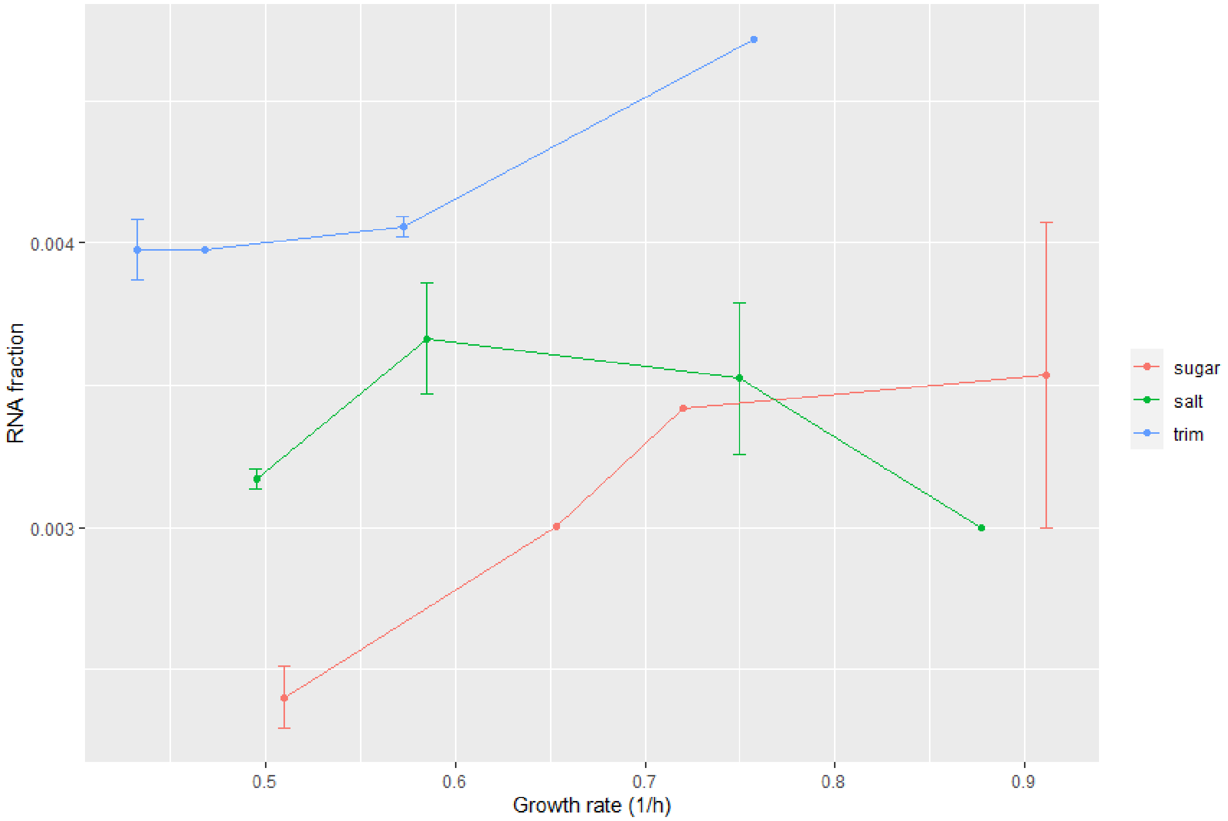


**Figure S13. Efflux pump expression is a small fraction of the S sector.** Efflux pump expression is ~1.5% the size of the S sector on glucose. Efflux pump genes comprise ~0.6% of S genes. The efflux pumps showed a dependence on conditions that cannot explain the present results. Their fraction decreases in poor sugars, increases for TMP, and remains relatively constant in NaCl.

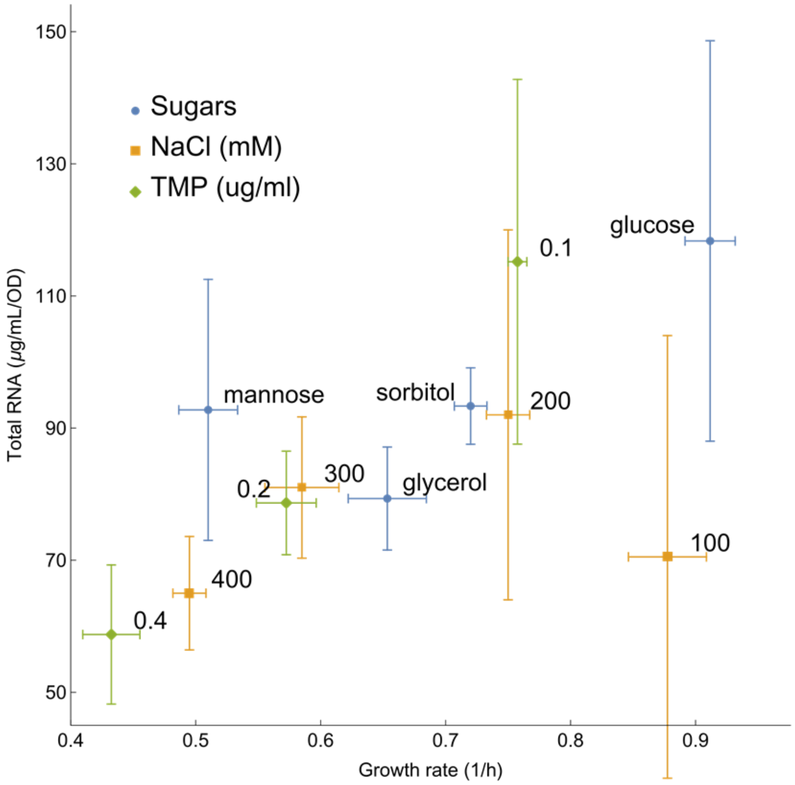


**Figure S14. Total RNA as in Fig 3F inset without normalization to glucose.** Error bars are standard error.

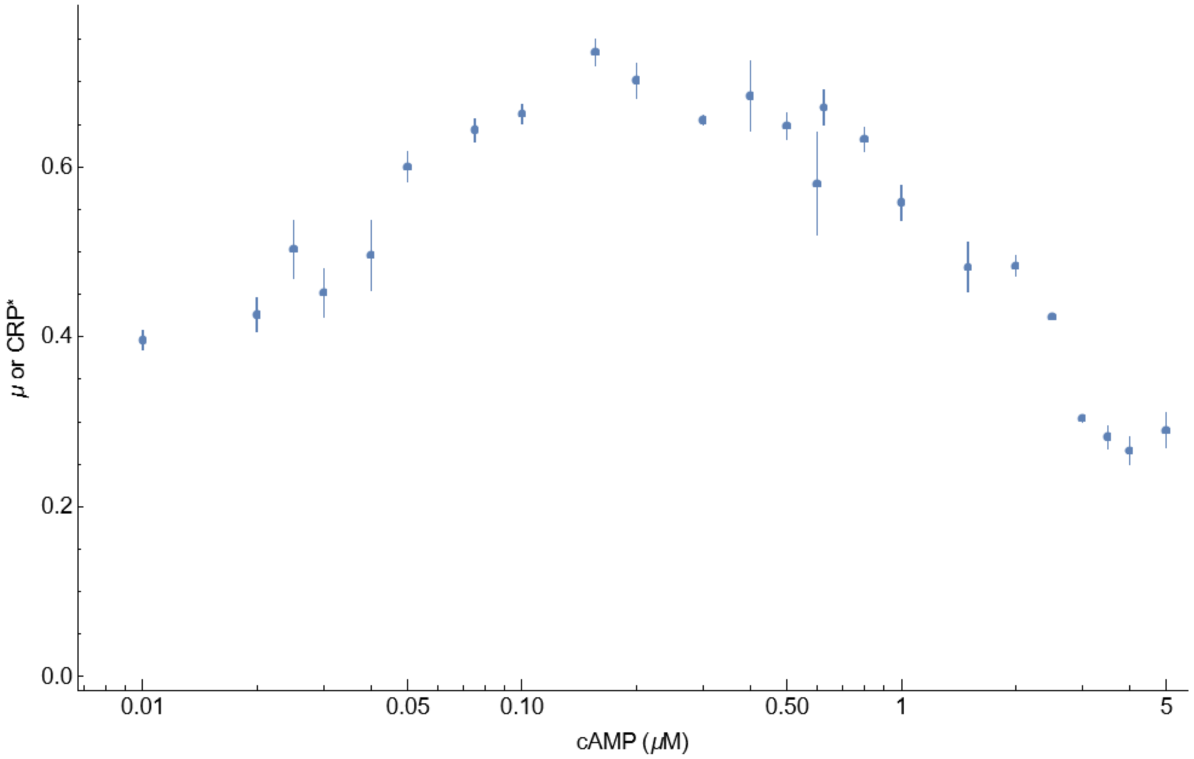


**Figure S15. Growth rate as a function of cAMP in MG1655ΔcyaAΔcpdA is a non-monotonic function.** The peak growth rate occurs at approximately 0.25 µM cAMP

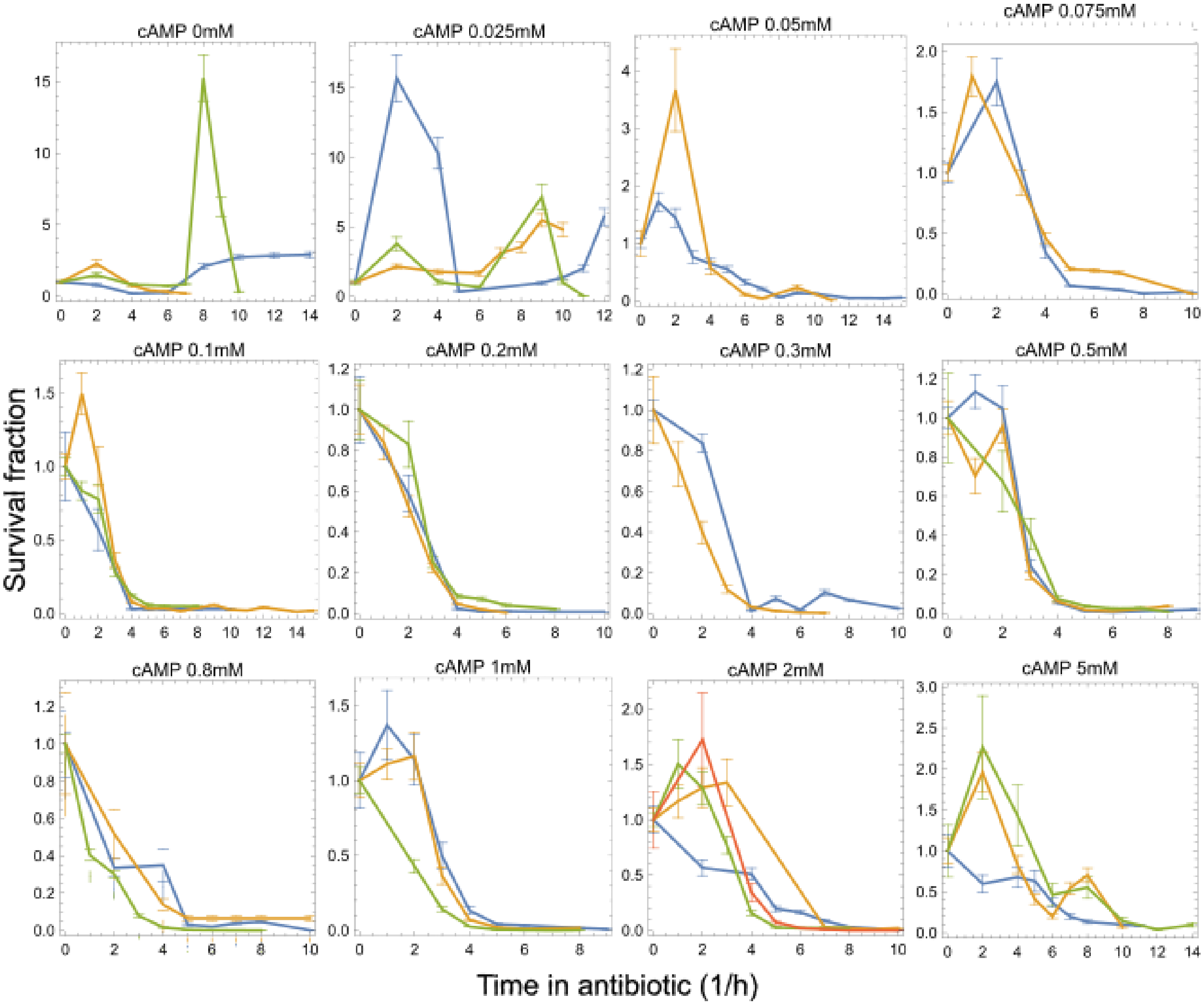


**Figure S16. All survival curves for MG1655 ΔcyaA ΔcpdA (U486) strain as determined by our automatic death assay.** Cultures were challenged with 15 µg/ml nalidixic acid. Different colors in each panel represent biological repeats. All conditions are minimal medium (M9) + glucose + indicated concentration of cAMP.

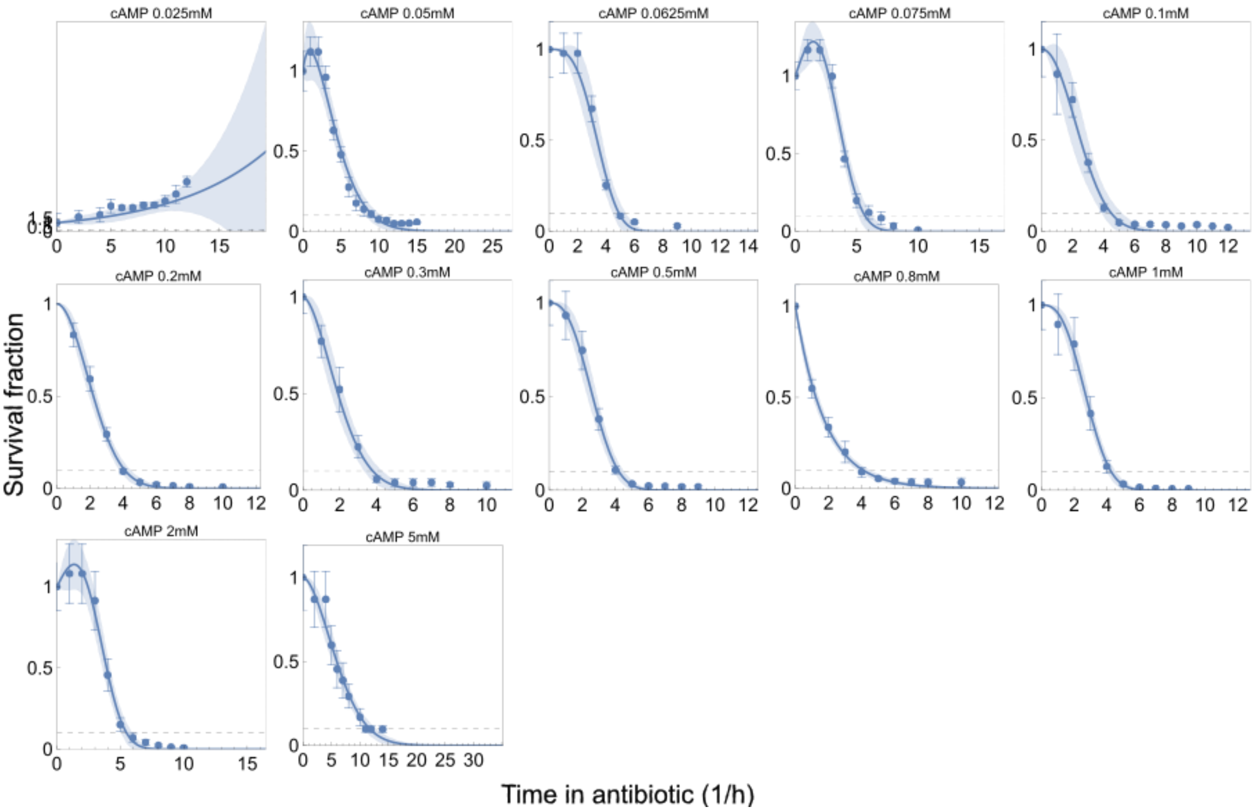


**Figure S17. Weibull death plus growth for averaged survival curves in U486 for presentation purposes.** Dashed line represents 10% survival. Shaded bands represent 95% confidence intervals on the fit. Points here are weighted averages of the data in Fig. S15. Death rates were calculated from similar fits to the individual replicates in Fig. S15 and then averaged.

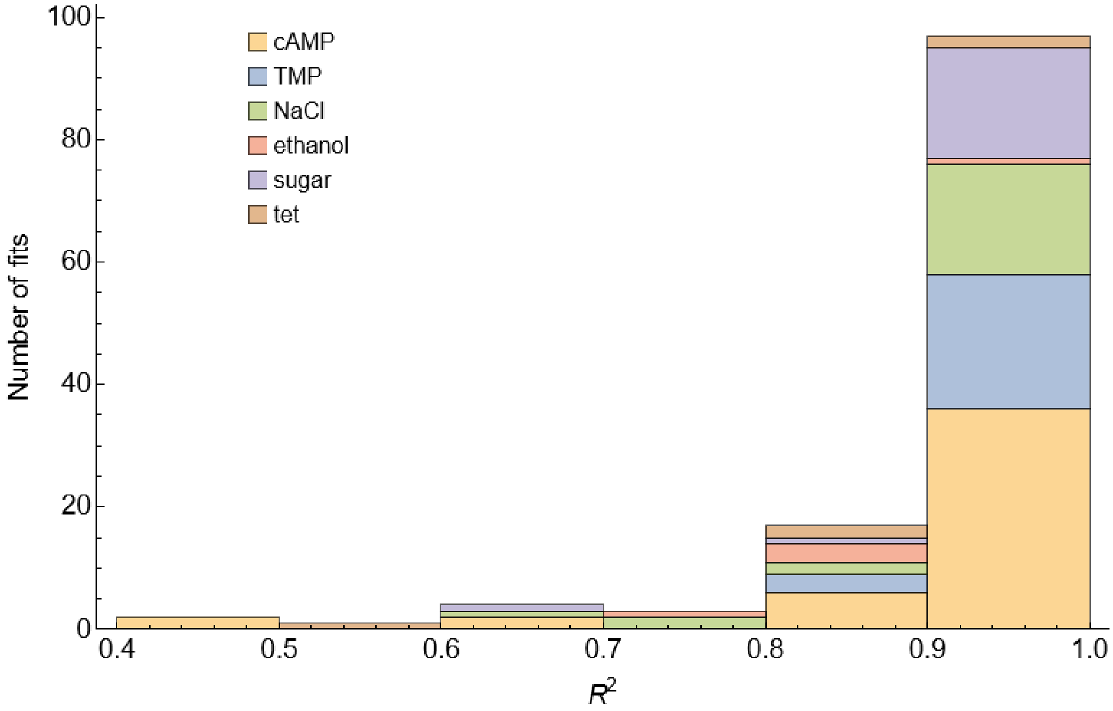


**Figure S18. $R^2$ for Weibull plus growth fits for all NCM3722 and U486 survival curves in Naladixic acid.** The $R^2$ values are taken from fits to data in Figs. S2 and S16.

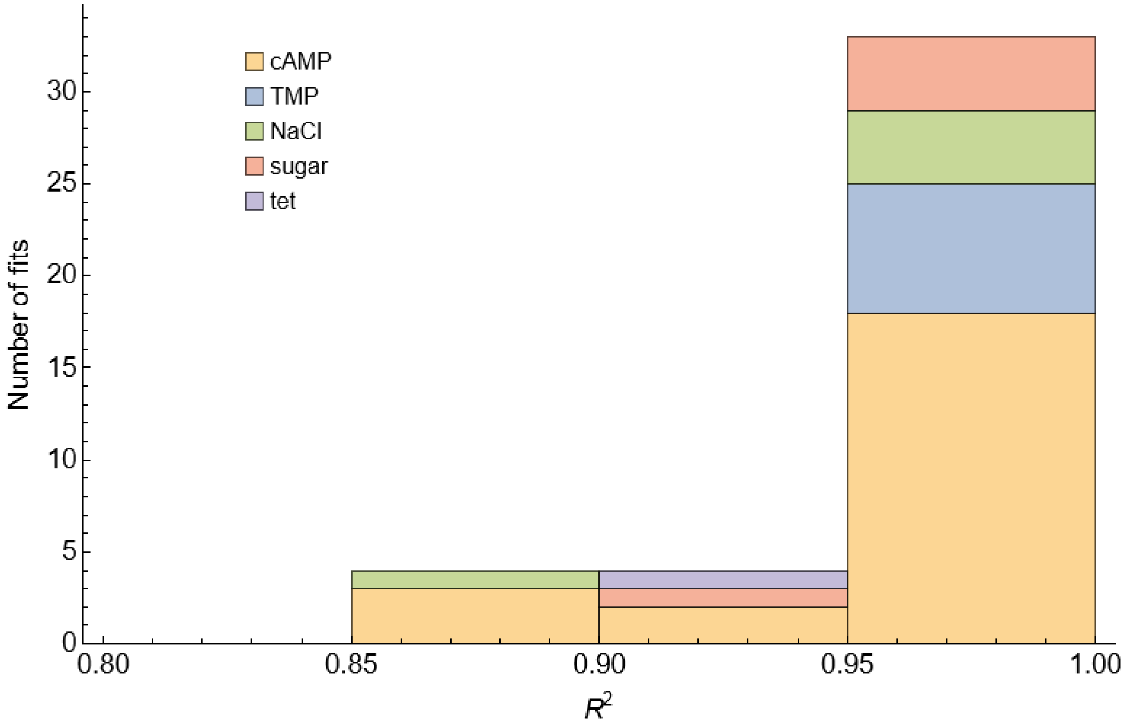


**Figure S19. $R^2$ for Weibull plus growth fits for all NCM3722 and U486 averaged survival curves in Naladixic acid.** The $R^2$ values are taken from fits to data in Figs. S3 and S17.

**Table S1. Parameter values for Fig. 3A (mean +/- standard error)**

| Parameter | Estimate |
|---|---|
| $\hat{\alpha}$ | 0.72±0.11 |
| $\hat{\beta}_{NaCl}$ | 1.41±0.14 |
| $\hat{\beta}_{TMP}$ | 0.88±0.13 |

**Table S2. Parameter values for Fig. 3D-F (mean +/- standard error)**

| Parameter | Estimate from sugar data | Estimate from NaCl data | Estimate from TMP data |
|---|---|---|---|
| $a$ | 1.619±0.002 | 6.97±0.06 | 7.22±0.04 |
| $S_G$ | 0.1165±0.0001 | 0.1167±0.0003 | 0.1360±0.0003 |
| $C_G$ | 0.2282±0.0002 | 0.2493±0.0002 | 0.2202±0.0002 |

**Table S3. Parameter values for Fig. 4A (mean +/- standard error)**

| Parameter | Estimate |
|---|---|
| $\hat{\alpha}$ | 1.20±0.05 |
| $\hat{\beta}$ | 2.63±0.42 |

**Table S4. Parameter values for Fig. 4B (mean +/- standard error)**

| Parameter | Estimate |
|---|---|
| $\mu_{PG}$ | 0.82±0.02 |
